# Long-term climatic stability drives accumulation and maintenance of divergent freshwater fish lineages in a temperate biodiversity hotspot

**DOI:** 10.1101/2023.03.08.531828

**Authors:** Sean James Buckley, Chris J. Brauer, Peter J. Unmack, Michael P. Hammer, Mark Adams, Stephen J. Beatty, David L. Morgan, Luciano B. Beheregaray

## Abstract

Anthropogenic climate change is forecast to drive regional climate disruption and instability across the globe. These impacts are likely to be exacerbated within biodiversity hotspots, both due to the greater potential for species loss but also to the possibility that endemic lineages might not have experienced significant climatic variation in the past, limiting their evolutionary potential to respond to rapid climate change. We assessed the role of climatic stability on the accumulation and persistence of lineages in an obligate freshwater fish group endemic in the southwest Western Australia (SWWA) biodiversity hotspot. Using 19,426 genomic (ddRAD-seq) markers and species distribution modelling, we explored the phylogeographic history of western (*Nannoperca vittata*) and little (*Nannoperca pygmaea*) pygmy perches, assessing population divergence and phylogenetic relationships, delimiting species and estimating changes in species distributions from the Pliocene to 2100. We identified two deep phylogroups comprising three divergent clusters, which showed no historical connectivity since the Pliocene. We conservatively suggest these represent three isolated species with additional intraspecific structure within one widespread species. All lineages showed long-term patterns of isolation and persistence owing to climatic stability but with significant range contractions likely under future climate change. Our results highlighted the role of climatic stability in allowing the persistence of isolated lineages in the SWWA. This biodiversity hotspot is under compounding threat from ongoing climate change and habitat modification, which may further threaten previously undetected cryptic diversity across the region.

## INTRODUCTION

Global biodiversity is increasingly threatened by anthropogenic climate change, with contemporary rates of extinction substantially higher than throughout much of geological history (Le Roux et al., 2019). These impacts are expected to be particularly exacerbated within regions that have experienced long-term climatic stability due to several convergent factors (Harrison & Noss, 2017). Firstly, species that have not been exposed to major climatic changes over their evolutionary histories are less likely to have evolved traits allowing adaptive responses to contemporary climate change (Sandoval-Castillo et al., 2020), including lower dispersal ability (Sandel et al., 2011) and thermal tolerance (Addo-Bediako, Chown, & Gaston, 2000) than species from more historically variable regions. Secondly, long-term climatic stability is often suggested to be a primary mechanism driving the spatial heterogeneity of biodiversity (Harrison & Noss, 2017; Sandel et al., 2011), particularly in hotspots of biodiversity (Habel et al., 2019; Carnaval, Hickerson, Haddad, Rodrigues & Moritz, 2009). Thus, understanding the role of climatic stability on biodiversity is important to predicting extinction risk under climate change.

The combination of conservation concerns within highly biodiverse regions is epitomised within global ‘biodiversity hotspots’. These regions are delineated by high species diversity, endemism, and the degree of habitat loss (>70% of primary vegetation; Myers, Mittermeier, Mittermeier, da Fonseca, & Kent, 2000). The disproportionate risk of extinction in biodiversity hotspots is also exacerbated by the fact that a considerable proportion of global biodiversity remains undocumented (Joppa, Roberts, Myers, & Pimm, 2011), including cryptic species (Adams, Raadik, Burridge & Georges, 2014; Struck et al., 2018). Accurately delimiting species remains a critical component of conservation management at both the taxon-specific (through revision of species classifications and associated legislation) and regional (through identifying hotspots of cryptic diversity) scales.

The first (of only two) biodiversity hotspots to be declared for Australia was the temperate southwest, commonly referred to as the Southwest Western Australia (SWWA) hotspot or the Southwest Australia Floristic Region (Hopper & Gioia, 2004; Myers et al., 2000). This region features high floristic diversity and endemism, with >8,000 species of plants recorded and >4,000 of those endemic to the region (Gioia & Hopper, 2017). While species diversity appears lower for animal groups (∼500 vertebrate species), relatively high rates of endemism (∼23%) suggest that this pattern extends to non-plant taxa as well (Rix et al., 2015). Despite the breadth of biodiversity, the SWWA features a simplistic landscape with limited topographic variation (Funnekotter, Millar, Krauss, & Nevill, 2019), no major river drainage divides, stable geology (Hopper & Gioia, 2004) and little climatic variation since the Pliocene (Spooner, De Deckker, Barrows, & Fifield, 2011). Within the region, biogeographic subdivisions have been delineated primarily based on rainfall, including the High Rainfall Province (HRP, >600mm annual rainfall) and the Transitional Rainfall Province (TRP, 300 – 600mm annual rainfall; Hopper & Gioia, 2004; Rix et al., 2015).

Several biogeographic mechanisms have been proposed to explain the high biodiversity of the SWWA (detailed in Rix et al., 2015). These include ancient Gondwanan lineages that diverged throughout the Mesozoic until the late Eocene ∼100 Ma (Hopper, Smith, Fay, Manning, & Chase, 2009); vicariantly-isolated mesic lineages that diverged from eastern Australian lineages during the Miocene (14–16 Ma; Buckley et al., 2018; Crisp & Cook, 2007; Rix & Harvey, 2012); as well as *in situ* diversification (Hopper & Gioia, 2004). Despite the apparent lack of topographic or environmental barriers (Cowling & Lombard, 2002), SWWA studies have demonstrated both interspecific (speciation) and intraspecific (phylogeographic structure) diversification primarily associated with late Miocene – early Pliocene aridification and contraction of mesic refugia (Byrne et al., 2011; Rix et al., 2015; Rix & Harvey, 2012). These disparate diversification histories suggest that the persistence of lineages is a key factor underlying the biodiversity of the SWWA.

The SWWA classification as a biodiversity hotspot is also driven by extensive habitat loss and deforestation, with ∼70% of native land vegetation cleared primarily for agricultural purposes (Habel et al., 2019; Monks et al., 2019). These threats are exacerbated by recent and rapid climatic changes, with a 10 – 15% decrease in rainfall since the 1970s (Ali et al., 2012) and a 1.1°C increase in temperature over the last century (Hallett et al., 2018). Anthropogenic climate change has already been implicated in regional population and fitness declines of plants (e.g., Monks et al., 2019) and is considered a key threatening process for threatened vertebrates (Stewart et al., 2018). Understanding the potential impacts of climate change in the SWWA requires a better understanding of the role of history climatic stability on species persistence, as well as better documentation of existing biodiversity.

Freshwater taxa are ideal models for investigating phylogeographic history as their limited dispersal capacity, reliance on constrained habitats, and high propensity for cryptic diversity (e.g., Buckley, Brauer, Unmack, Hammer, & Beheregaray, 2021; Waters, Burridge, & Craw, 2020) make them effective indicators of historical environmental change. Within the SWWA, the western pygmy perch (*Nannoperca vittata*) and little pygmy perch (*Nannoperca pygmaea*) demonstrate these traits, given their late Miocene origins (Buckley et al., 2018), limited dispersal capacity (Beatty, Morgan, Rashnavadi, & Lymbery, 2011), and role as ecological specialists (Allen, Morgan, Close, & Beatty, 2020). *Nannoperca vittata* is significantly more widespread and occurs throughout the HRP, whereas *N. pygmaea* has a restricted occurrence within only three rivers and a lake in the south-eastern HRP (Allen et al., 2020). While both species show some similar ecological characteristics such as body plan and reproductive strategy, they show marked differences in body size, morphology, growth rate, salinity tolerance and reproductive timing (Allen et al., 2020). Additionally, clear-cut genetic differentiation in allozymes (Morgan, Beatty, & Adams, 2013), mitochondrial genes (Unmack, Hammer, Adams, & Dowling, 2011) and genome-wide markers (Buckley et al., 2018), and a lack of observed hybridisation with *N. vittata* (Morgan et al., 2013; Allen et al., 2020), corroborates the identity of *N. pygmaea* as a distinct species. This is reflected by their strong evolutionary distinctiveness, with divergence between the two species estimated at ∼4 million years ago (Buckley et al., 2018; Unmack et al., 2011). In addition, up to three distinct cryptic species have been suggested within *N. vittata* based on allozyme, mitochondrial and nuclear data (Unmack et al., 2011), and genome-wide data (Buckley et al., 2018). However, these studies were either lacking in genetic resolution (Unmack et al., 2011) or geographic sampling (Buckley et al., 2018). Additionally, *Nannoperca pygmaea* is currently listed as Endangered under both national and state legislation, whilst *N. vittata* is unlisted with no local legislative protections (Allen et al., 2020). Thus, a thorough investigation of phylogeographic patterns across the region is required to refine hypotheses of species delineation and inform conservation management for these species.

Here, we investigate the role of long-term climatic stability in the SWWA biodiversity hotspot on the diversification and accumulation of lineages in a group of freshwater fishes. We used genomic (ddRAD-seq) data to characterise divergence across the clade based on genetic differentiation, phylogenetic patterns, and species delimitation. Additionally, we used species distribution modelling to reconstruct species and lineage distributions since the Pliocene and predict distribution changes under future climate change. We hypothesised that long-term climatic stability across the biodiversity hotspot would lead to both the maintenance of divergent and isolated genetic lineages, including cryptic species, and long-term stable species distributions. Conversely, we predicted that future climate change in the SWWA poses a significant conservation threat for these species, evidenced by projected range contractions.

## METHODS

### Sample Collection and Library Preparation

Sampling sites were selected to capture the full distribution of all lineages, including a disjunct and potentially relictual population found in the northernmost extreme of the TRP (Fig. 1; Buckley et al., 2018; Unmack et al., 2011). We sampled a total of 25 *N. vittata* from seven sites and eight *N. pygmaea* from two sites (Supplementary Table S1): although sparse, this sampling design spans the full diversity of *Nannoperca* in the region, including all described and previously suggested cryptic species within a broader *N. vittata* species complex (Buckley et al., 2018; Unmack et al. 2011) and all intraspecific lineages (Adams et al., unpublished allozyme data). An additional four Yarra pygmy perch (*Nannoperca obscura*) were included as an outgroup for phylogenetic analyses (Buckley et al., 2018). Specimens were collected using electrofishing, dip-, fyke- or seine-netting (see Allen et al., 2020). Either the caudal fin or the entire specimen was stored at −80°C at the South Australian Museum, or in 99% ethanol at the Molecular Ecology Lab at Flinders University.

**Fig. 1.**
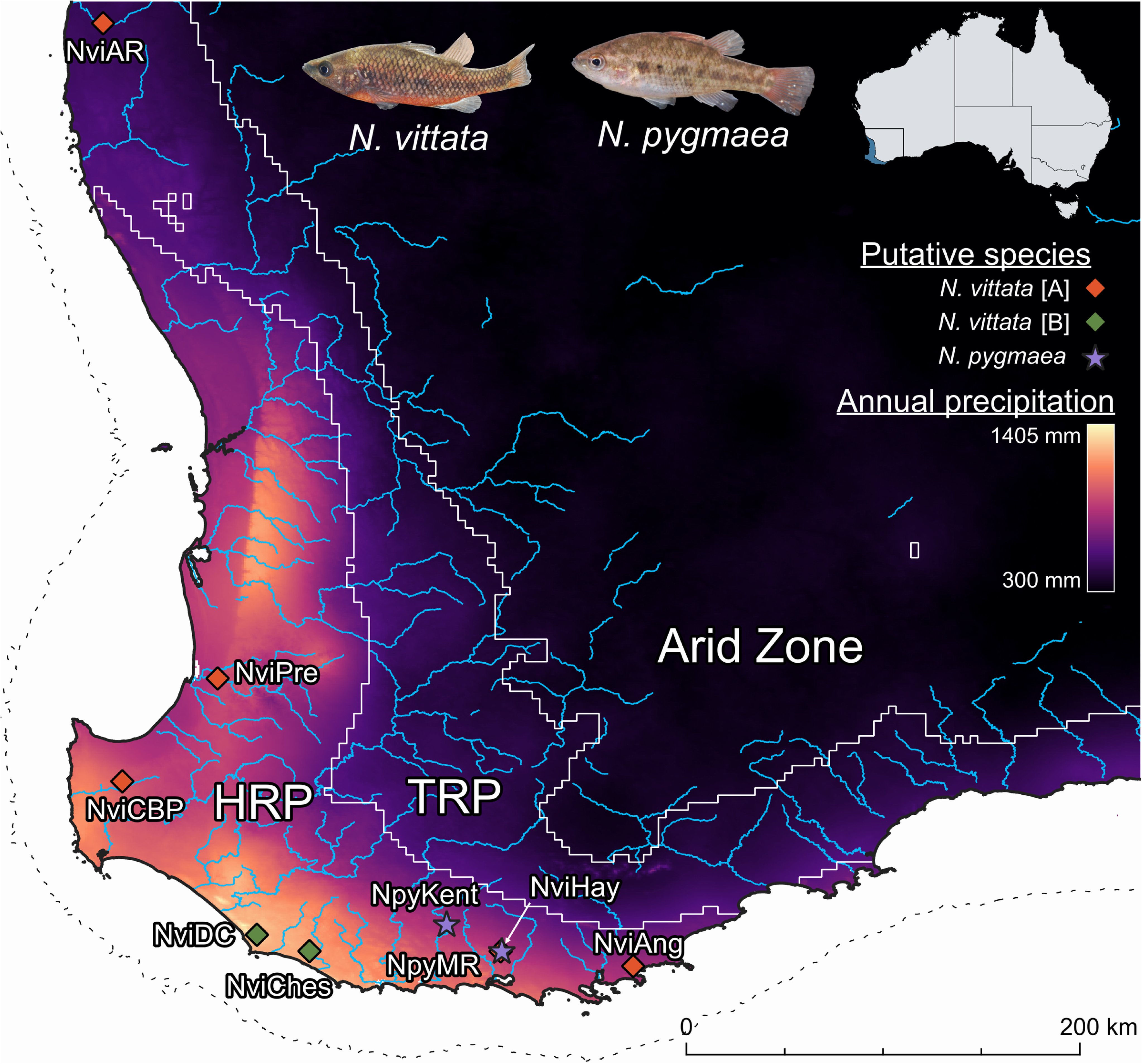
Map of sampling sites used within this study, and major rivers across the SWWA. Localities for *Nannoperca vittata* are indicated by diamonds and localities for *Nannoperca pygmaea* are indicated with stars. Locality colours indicate putative species (see Results). Colouration of the map indicates annual precipitation (derived from WorldClim 2.1). Solid white lines indicate biogeographic province boundaries and the dashed line indicates the maximum landmass extent exposed during the Last Glacial Maximum (LGM). Inset map indicates extent of distribution to the Australian continent. Fish images demonstrate described (morphological) species and current taxonomy. Photos courtesy of Stephen Beatty. TRP = Transitional Rainfall Province. HRP = High Rainfall Province.

DNA was extracted from muscle tissue or fin clips using a combination of a modified salting-out method (Sunnucks & Hales, 1996) and a Qiagen DNeasy kit (Qiagen Inc., Valencia, CA, USA). Genomic DNA was checked for quality using a spectrophotometer (NanoDrop, Thermo Scientific), integrity using 2% agarose gels, and quantity using a fluorometer (Qubit, Life Technologies). The ddRAD genomic libraries were prepared in house following Brauer, Hammer, & Beheregaray (2016). Of the 33 samples, eight were paired-end sequenced on an Illumina HiSeq 2000 at Genome Quebec (Montreal, Canada) as part of a previous phylogenomic study (Buckley et al., 2018). The remaining 25 samples were single-end sequenced on a single lane of Illumina HiSeq 2500 at the South Australia Health and Medical Research Institute in Adelaide.

### Filtering and Alignment

Sequences were demultiplexed using the ‘process_radtags’ module of Stacks 1.29 (Catchen, Hohenlohe, Bassham, Amores, & Cresko, 2013), allowing up to 2 mismatches in the 6 bp barcodes. Barcodes were removed and sequences trimmed to 80 bp to remove low-quality bases from the end of the reads. Cut reads (forward only for paired-end) were aligned using PyRAD 3.0.6 (Eaton, 2014), and further cleaned by removing reads with >5 bp with a Phred score < 20. Loci were retained if they occurred in at least ∼80% of samples (30) within the dataset. For SNP-based analyses, a single SNP per ddRAD locus was subsampled to reduce the impact of linkage disequilibrium.

### Population Divergence and Population Clustering

We first assessed population divergence by estimating mean pairwise uncorrected genetic distances (*p*-distances) between populations using PAUP* 4 (Swofford, 2002) based on the full sequences of all ddRAD loci. Population clustering was assessed using a PCoA of all SNPs in dartR (Gruber, Unmack, Berry, & Georges, 2018). We also calculated the number of fixed differences between pairwise population comparisons (Unmack et al., 2022), with SNPs considered fixed differences at a threshold of 0.05 (i.e. >95% frequency in one population and <5% in the other) to account for sequencing errors or ‘near fixation’ (Gruber et al., 2018).

### Phylogenetic Analysis

A maximum likelihood (ML) phylogeny was estimated using RAxML 8.2.11 (Stamatakis 2014) and the concatenated sequence alignment to determine evolutionary relationships and guide species delimitation analyses. The ML phylogeny was estimated under the GTR+Γ model of evolution and 1,000 resampling estimated log-likelihood bootstraps. A ML phylogeny with the alignment partitioned by ddRAD locus was also estimated, as well as estimating individual gene trees per locus, using IQ-TREE2 (Minh et al., 2020b). This was done to account for the potential impact of genome-wide heterogeneity in evolutionary rates (Liu, Xi, & Davis, 2015). Concordance between gene trees and the partitioned phylogeny was estimated using site and gene concordance factors (Minh, Hahn, & Lanfear, 2020a), with a summary species tree estimated using ASTRAL-III (Zhang, Rabiee, Sayyari, & Mirarab, 2018) and assuming each population as an individual ‘species’.

### Species Delimitation and Divergence Time Estimates

The species tree and delimitation were estimated using SNAPP 1.5.0 (Bryant, Bouckaert, Felsenstein, Rosenberg, & RoyChoudhury, 2012) within BEAST 2.6.1 (Bouckaert et al., 2019). We iteratively tested nine different scenarios of species composition, ranging from two to nine species (i.e. each population as a separate species) based on phylogenetic patterns (see Fig. 2d). Given that SNAPP can resolve species identities with only a few thousand SNPs (Leaché, Fujita, Minin, & Bouckaert, 2014), and to reduce computational time, we subsampled the alignment down to the two individuals with the lowest missing data per population and 5,000 randomly selected SNPs. We used broad priors with gamma distributions for speciation rate (λ; α = 3, β = 2.5) and population sizes (θ; α = 2.85, β = 955.27) based on sequence divergence for all scenarios to cover possible parameter ranges. Mutation rate priors were left at their default settings. Two separate chains were run per model to assess convergence of parameter estimates. Models were run for at least 10 million generations and/or until ESS > 200 was reliably achieved. Model traces were visualised using Tracer 1.5 (Rambaut & Drummond, 2009) with the first two million generations discarded as burn-in. Species composition likelihoods were estimated using an AIC through Markov chain Monte Carlo analysis (AICM; Raftery, Newton, Satagopan, & Krivitsky, 2006). Although other methods such as reverse-jump MCMC via path-sampling or stepping-stone analysis are more widely used (Grummer, Bryson, & Reeder, 2013), given the large size of the dataset we opted to use AICM based on its reasonable performance and reduced computational demand (Baele et al., 2012).

**Fig. 2.**
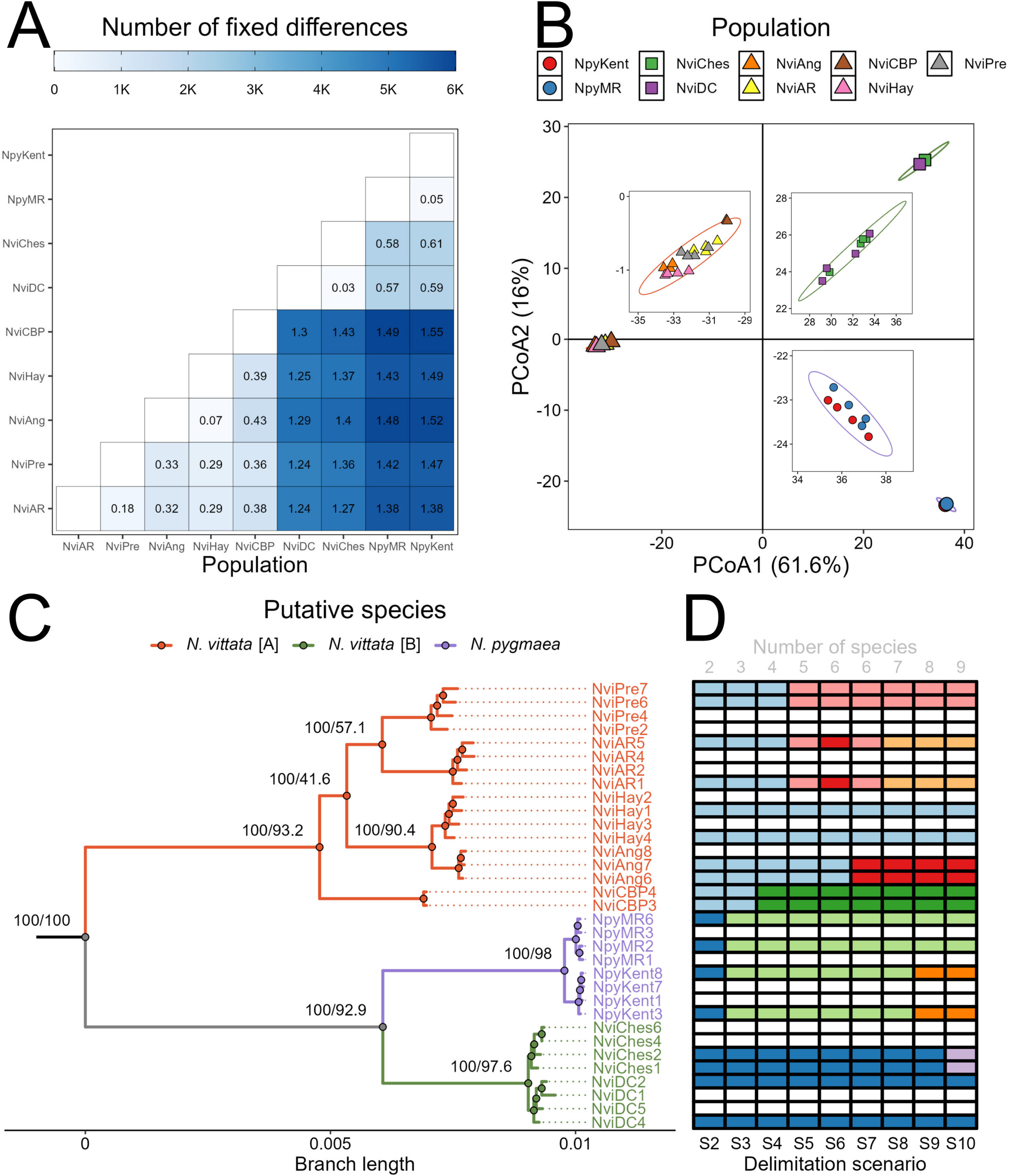
Population divergence and clustering. **A** Heatmap of pairwise population summaries of divergence. Numbers within cells indicate pairwise uncorrelated genetic distances (p-distance; %) based on 19,426 ddRAD loci, whilst colours indicate the number of fixed differences within the 18,177 unlinked SNPs. **B** PcoA of 18,177 unlinked SNPs. Each point represents the centroid per population with shapes indicating assignment to a putative species (circles = *Nannoperca pygmaea*, triangles = *Nannoperca vittata* [A] and squares = *N. vittata* [B]). Insets depict close-ups of individuals within each cluster. **C** Maximum likelihood tree estimated using 19,426 concatenated ddRAD loci and IQ-TREE2. Nodes are labelled with bootstrap support estimated using 1,000 RELL bootstraps and site concordance factors (respectively). Branches are coloured by putative species. **D** Allocation of samples to species delimitation scenarios in SNAPP. Each column represents a single model, with each row corresponding to the aligned sample in the phylogenetic tree. Cell colours indicate allocation to species, with cells of the same colour indicating one species. Blank rows indicate samples that were not used in species delimitation.

Divergence times across the complex were inferred by using an extension of SNAPP described in Stange, Sanchez-Villagra, Salzburger, & Matschiner (2018). We calibrated the oldest divergence in the tree using a normal distribution with mean of 9.27 Ma and standard deviation of 0.51 (95% CI = mean ± 1 Ma) based on previous divergence estimates for all pygmy perches (Buckley et al., 2018). We applied broader ranges around calibration nodes than previously suggested to accommodate potential variation in calibration age not captured by the methods in that study. The topology of the tree was fixed using the ML tree and the model run for one million generations. Confidence intervals of divergence times were inferred using TreeAnnotator 2.6 (Drummond & Rambaut, 2019) and 95% posterior probabilities.

### Historical Admixture and Introgression

We tested whether populations and lineages were historically isolated using two different allele frequency-based approaches. First, we determined historical connectivity across populations based on the SNP dataset using TreeMix (Pickrell & Pritchard, 2012). The number of migrations within the model were iteratively increased from none to nine, with the fit of each model estimated using the covariance matrices and overall tree likelihood. Additionally, we calculated the percentage of variation explained per migration model (https://github.com/wlz0726/Population_Genomics_Scripts/tree/master/03.treemix), with the best supported number of migrations determined by the asymptote of likelihood. We also determined whether substantial introgression occurred between putative species (see Results) suggested by species delimitation approaches using an ABBA-BABA test (Martin, Davey, & Jiggins, 2014) in D-Suite (Malinsky, Matschiner & Svardal, 2021). Introgression was determined assuming a pattern of divergence following the phylogenetic tree and summarised based on Patterson’s D (Patterson et al., 2012) and the fraction of introgressed alleles (f_4_-ratio).

### Species and Lineage Distribution modelling

Species distribution models (SDMs) for *N. vittata* were estimated using an ensemble modelling approach within biomod2 (Thuiller, Lafourcade, Engler, & Araújo, 2009). SDMs were projected from contemporary conditions to the Pliocene using the PaleoClim database (Brown, Hill, Dolan, Carnaval, & Haywood, 2018), including the Late Holocene (4.2–0.3 Kya), Mid Holocene (8.326–4.2 Kya), Early Holocene (11.7– 8.326 Kya), Younger Dryas Stadial (12.9–11.7 Kya), Bølling-Allerød (14.7–12.9 Kya), Heinrich Stadial 1 (17.0–14.7 Kya), Last Interglacial (∼130 Kya), MIS19 (∼787 Kya), mid Pliocene warm period (3.205 Ma), and M2 (∼3.3 Ma) phases. Additionally, we extrapolated SDMs to future conditions (2020 – 2040, 2040 – 2060, 2060 – 2080 and 2080 – 2100) under three climate change scenarios (Shared Socioeconomic Pathways, SSPs: SSP126, SSP245 and SSP585) using the ACCESS-CM2 global circulation model (Bi *et al*. 2020) derived from CMIP6 via WorldClim (Fick & Hijmans, 2017). We opted for this model based on its performance in the SWWA environment, with low error and high skill score (Moise et al. 2015).

Occurrence records for all *N. vittata* were obtained from a combination of sampled sites within this and past studies (Allen et al., 2020; Buckley et al., 2018; Morgan, Gill, & Potter, 1998; Morgan et al., 2013; Unmack et al., 2011), as well as from the Atlas of Living Australia (http://www.ala.org.au). To reduce the impact of spatial autocorrelation (Elith et al., 2011), we sampled a single occurrence per environmental raster cell. We further filtered the occurrence data by removing occurrences outside the known distribution of species (P. Unmack, pers. comm.; Allen *et al*. 2020) to remove potentially erroneous records. This resulted in a final dataset of 114 observations used within the SDMs.

Highly correlated climatic variables (*r* < |0.8|) were pruned from the dataset based on a Pearson’s correlation test within SDMToolbox (Brown, Bennett, & French, 2017), resulting in eight climatic layers used across the models (Table S2). The final input variables were annual mean temperature (bio1), isothermality (bio3), mean temperature of the wettest quarter (bio8), mean temperature of the driest quarter (bio9), mean temperature of coldest quarter (bio11), annual precipitation (bio12), precipitation of the driest month (bio14), precipitation seasonality (bio15) and sea-level corrected elevation. For the three oldest time periods, bio3 was unavailable and thus not included. We generated three replicates of 1,000 pseudoabsences randomly from the background >30 km from occurrences to reduce the likelihood of generating false absences within habitable areas. Each dataset was replicated three times, with 70% of sites independently and randomly subset to train the model (*n* = nine datasets).

The SDMs were estimated using four separate algorithms: classification tree analysis (CTA), multiple adaptive regression splines (MARS), maximum entropy (MaxEnt) and random forest (RF) (*n* = 36 models total). These algorithms cover a range of different statistical approaches (Hao, Elith, Guillera-Arroita, & Lahoz-Monfort, 2019), and model averaging in an ensemble framework is expected to reduce bias associated with any single method (Marmion, Parviainen, Luoto, Heikkinen, & Thuiller, 2009). Each model was evaluated using the relative operating characteristic (ROC) and the true skill statistic (TSS). Individual algorithms were checked for consistency by averaging models per method under contemporary conditions. Ensemble SDMs amalgamating the results of all SDMs per time period were generated, excluding models with TSS < 0.7. We also converted all individual and ensemble models into binary presence-absence maps based on the TSS within biomod2 and calculated the distribution area.

We also estimated individual lineage-specific distribution models (LDMs) for each putative *N. vittata* species by assigning each of the 114 *N. vittata* observations to the nearest sampled lineage (where putative species could be confirmed), removing any observations >50km (half the distance between the two nearest points across putative species). This resulted in 46 and 30 *N. vittata* [A] and [B] observations, respectively. All LDMs were estimated under the same parameters and algorithms as the SDMs, albeit pseudoabsences for *N. vittata* [A] were generated at >85km from presences (to prevent pseudoabsences occurring in *N. vittata* [B] habitat). All models were projected back to the Last Interglacial and across all future climate change conditions.

## RESULTS

### Bioinformatics

The combined sequencing runs returned a total of 71.44 million reads, with an average of 2.16M reads per sample. After quality control and alignment of sequences, a dataset of 19,426 ddRAD loci was obtained across all putative species. This alignment contained 18,177 putatively unlinked biallelic SNPs and an average of 4.93 (±1.76)% missing data per sample (Supplementary Fig. S1).

### Population Divergence

Pairwise genetic distances and fixed differences inferred highly divergent structure across the species complex. Pairwise population averages of genetic differences ranged from 0.03% to 1.55% whist the number of pairwise fixed differences ranged from 8 to 5,947 SNPs (Fig. 2a). A PCoA demonstrated three major and highly divergent clusters, with the two primary axes explaining a combined total of 77.6% of the variation (Fig. 2b). The three clusters are herein referred to as *N. pygmaea, N. vittata* [A] (widespread across SWWA; 5 sites) and *N. vittata* [B] (restricted to south-central HRP; 2 sites). Similar clusters could be observed within the pairwise *p*-distances (>0.5% between clusters) and by the number of fixed differences (>2000 between clusters).

### Phylogenetic Analysis

Partitioning the alignment by ddRAD loci did not affect the topology or branch lengths of the tree, and site concordance factors supported this tree (Supplementary Fig. S2a–S2c). The summary species tree likewise showed strong support for the separation of lineages, with posterior probabilities of one for almost all nodes (Supplementary Fig. S2d). The highly divergent nature of lineages across the group was supported by the concatenated phylogeny, with all population-level and above nodes supported by 100% bootstrap values (Fig. 2c). The phylogeny demonstrated three divergent lineages matching the clusters defined earlier. Despite their co-occurrence, the sympatric populations of *N. vittata* [A] and *N. pygmaea* (NviHay and NpyMR) were not closely associated within the phylogenetic tree; instead, *N. vittata* [B] was the sister lineage to *N. pygmaea*.

### Species Delimitation and Divergence Time

Species delimitation models increased in likelihood with an increasing number of species, with the highest likelihood and lowest AICM for a delimitation scenario considering all populations as separate species (Fig. 2d; scenario S10). However, there was a noticeable plateau in AICM values after S5, with greater numbers of species conferring relatively small increases in model likelihood beyond this point (Supplementary Fig. S3). Divergence time estimates suggested that most lineages diverged from one another between 1 and 5 Ma, with most populations diverging more recently, within the last 300 Kya (Fig. 3a). For all nodes, there was minimal overlap in 95% confidence intervals.

**Fig. 3.**
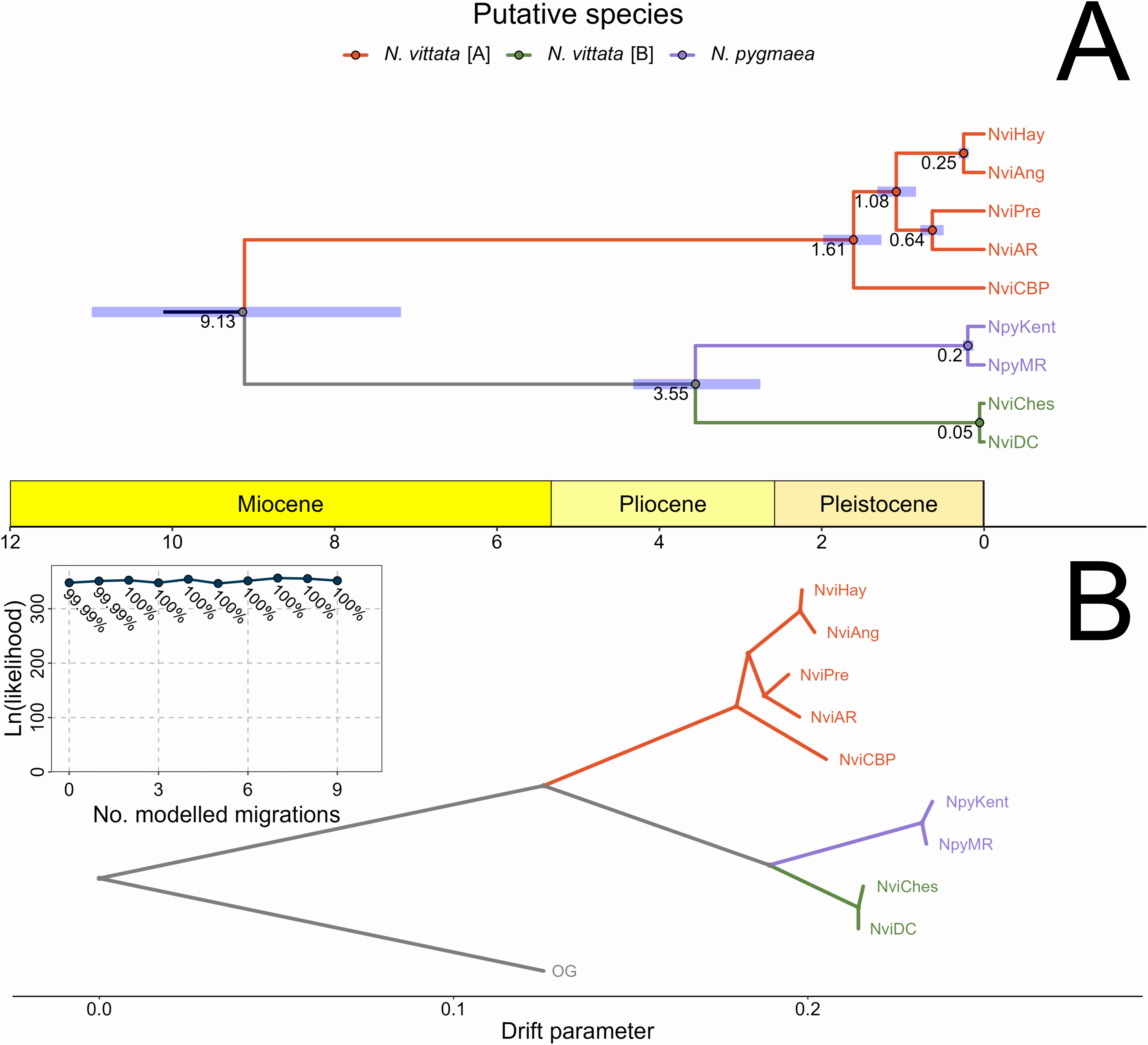
Divergence times and historical migration patterns. **A** Chronogram of divergence time estimates across the species complex using SNAPP. Node labels indicate median divergence times, and error bars show 95% posterior probabilities from 1M simulations. Lineages are coloured according to putative species. **B** Best supported historical migration model using TreeMix, with no modelled migrations and lineages coloured by putative species. Inset depicts likelihood and percentage of variation explained (labels) per model under increasing number of migrations. OG = outgroup.

**Fig. 4.**
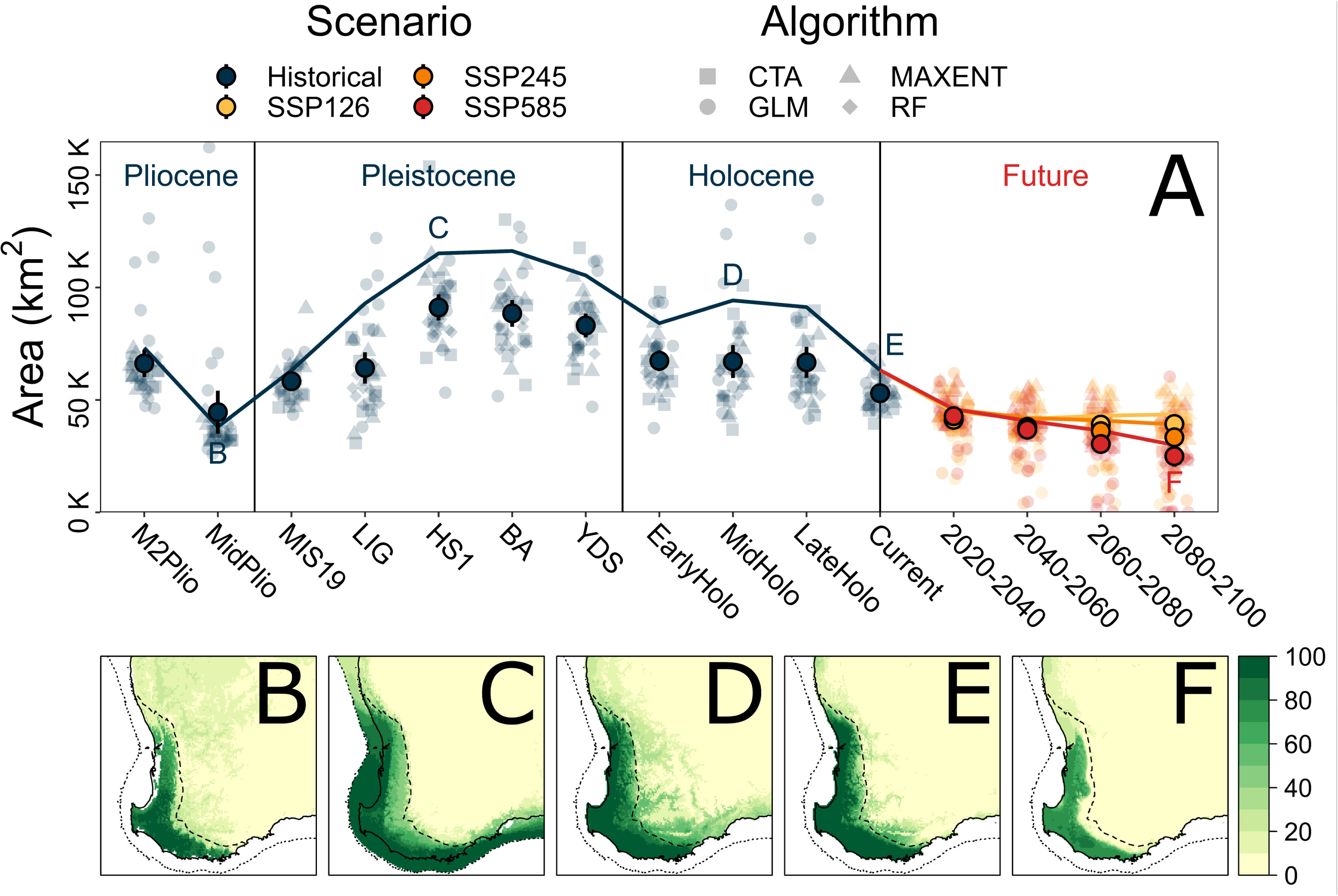
**Species distribution modelling for the *Nannoperca vittata* species complex**. **A** Estimated area of suitable habitat using binary distribution models, from oldest (left) to future (right). Solid line indicates estimates from ensemble (weighted mean) models, lighter points indicate estimates from individual models (*n* = 36; shapes indicate algorithm used) and solid points indicate mean estimates across all individual models (with standard error bars). Text labels indicate ensemble models represented in the bottom row. **B** Ensemble SDM for the mid-Pliocene warm period (3.205 Mya). **C** Ensemble SDM for HS1 (17 – 14.7 Kya). **D** Ensemble SDM for the mid-Holocene (8.326 – 4.2 Kya). **E** Ensemble SDM under contemporary conditions (1970 – 2000). **F** Ensemble SDM under SSP585 for 2080 – 2100.

### Gene Flow and Introgression

No signal of historical population connectivity was inferred across the clade, with TreeMix best supporting a tree that contained no migrations over models with any (Fig. 3b). This non-migratory model explained 99.99% of the variation in allele frequencies (Fig. 3b; Supplementary Fig. S4). Similarly, D-Suite results showed negligible evidence of introgression across putative species, with a Patterson’s *D* of 0.20 and f_4_-ratio of 0.028.

### Species Distribution Modelling

All models demonstrated high fit to the data, accurately capturing the full distribution of the species complex (Supplementary Fig. S5a). Ensemble models built separately for each method demonstrated highly similar areas of suitability, suggesting that variation among methods was minimal (Supplementary Fig. S5c). Annual precipitation (bio12) was the strongest driving variable across all models, with approximately double the variable importance of all others (Supplementary Fig. S5b). The ensemble projections of the SDMs over time showed little fluctuation in distribution extent since the Pliocene (Fig. 5a; Supplementary Figure S6). Across all time periods, the most significant portion of suitable habitat tended towards the coastal edges of the region, with some expansion across the continental shelf during lower sea levels but not inland (Fig. 5b – f). Future projections under all climate change scenarios indicated a range contraction would be likely by 2060, particularly within the northern and inland edges of the current distribution (Supplementary Fig. S7). More severe climate change scenarios (e.g., SSP585) are likely to result in an increasingly extensive range contraction and an overall reduction in habitat suitability, beyond that of historical projections.

Similar to the SDMs, historical LDMs appeared stable over time, with coastward expansions during lower sea levels (Supplementary Fig. S8 – S9). Future projections of LDMs also suggested range contractions under climate change, with *N. vittata* [A] predicted to have substantial northern and inland contractions under more severe climate change scenarios (Supplementary Fig. S10). Range contractions for *N. vittata* [B] were predicted to be even more extreme, leading to a total loss of climatic niche by 2100 under SSP245 and by 2080 under SSP585 (Supplementary Fig. 11).

## DISCUSSION

Several divergent lineages were identified across a species complex of freshwater fish endemic to a temperate biodiversity hotspot, including at least one cryptic species of pygmy perch. We demonstrated that these lineages probably persisted in isolation since the late Miocene (9–3.5 Ma) over a climatically stable region. Current and projected climatic conditions disproportionately threaten this region because its high lineage diversity appears to have evolved in the absence of major climatic and geological changes.

### Maintenance of Highly Divergent Lineages Through Climatic Stability

The highly divergent and ancient nature of lineages within the clade was likely facilitated by the long-term climatic stability of the region, with divergence events spanning the Miocene and Pliocene. Although major environmental changes associated with aridification during the Pliocene impacted the biota of the SWWA (Hopper & Gioia, 2004), temperature and rainfall remained relatively constant throughout the glacial cycles of the Pleistocene, especially within the HRP (Rix et al., 2015). The stable species distribution reflected these relatively constant climatic conditions since the Pliocene (Spooner et al., 2011), with a coastward expansion when sea levels were lower during glacial maxima. Similar patterns of limited variation in inland regions have been shown for several coastal plant species in the SWWA (Nevill, Bradbury, Williams, Tomlinson, & Krauss, 2014; Nistelberger et al., 2014), suggesting that distribution extent was consistently limited by aridity and temperature in inland habitats (Brouwers et al., 2012). These spatial patterns reflect the intensity of aridification further inland compared to the relatively benign HRP (Rix et al., 2015), and likely placed significant limitations on inland range expansions of *N. vittata*.

While causation of divergences within the clade was not inferred here, the historical demographic isolation of lineages, the overall low genetic diversity and low dispersal capacity of pygmy perches (Brauer et al., 2016; Buckley et al., 2018) together suggest that their persistence must owe, in part, to an enduring stable habitat. Overall, these results add to a growing body of evidence that broad-scale and long-term climatic stability in the SWWA has driven the accumulation of regional biodiversity (Supplementary Table S3). These endemic species encompass an array of lineages ranging from Gondwanan relicts originating as early as the Cretaceous to late Miocene – early Pliocene diversification events (Rix et al., 2015), reflecting both ancient and recent lineages (Sundaram et al., 2019). These concerted patterns thus demonstrate how biodiversity in SWWA has persisted and accumulated over millions of years due to the region’s climatic stability.

### Diversification of Western Australian Pygmy Perches

Our results highlight a hierarchy of divergence across Western Australia’s *Nannoperca* clade, ranging from morphologically differentiated and cryptic species to divergent populations. No historical migration nor introgression was detected across the clade, indicating that the inferred evolutionary distinctiveness is a product of long-term isolation. Summaries of population divergence largely identified three major clusters, including two separate *N. vittata* lineages, one of which was sister to *pygmaea*. This was supported with phylogenetic analysis, which highlighted the paraphyletic nature of the species name *N. vittata* (although *N. pymgaea* was also recently split from the *N. vittata* complex).

Species delimitation results implied the presence of a higher number of putative species within the complex, with greatest support for a model considering each population a separate species, and an approximate plateau in likelihood at five species. However, multispecies coalescent approaches such as SNAPP have been demonstrated to over-split lineages (Chambers & Hillis, 2020), particularly when populations are highly structured and divergent (Derkarabetian, Castillo, Koo, Ovchinnikov, & Hedin, 2019). Thus, we instead suggest that the genomic data unequivocally indicate that at least three species are present in the clade corresponding to the lineages *N. pygmaea*, *N. vittata* [A] and *N. vittata* [B], following similar nomenclature in Unmack et al., 2011. This conservative interpretation of three species both resolves the current paraphyly of *N. vittata* and avoids the risk of over-splitting (Coates, Byrne, & Moritz, 2018). A morphological revision is required to diagnose the three lineages (i.e. especially *N. vittata* [B]), and then match these to the morphotypes of available names of which two are potentially available; *N. vittata* [non-specific type locality] and *Nannoperca viridis* [King George Sound near Albany]. It is more likely that type material of *N. vittata* s.s. was collected from within the wide range of *N. vittata* [A] compared to the more inaccessible south-central HRP (habitat of *N. vittata* [B]), and if so *N. viridis* occurring well to the east is likely to remain in synonymy with *N. vittata* s.s., and a new description required for *N. vittata* [B].

Taxonomic revision of the species complex is critical given that species are the most common operational unit of conservation management (Stanton et al., 2019), and particularly, are the focus of protective legislation (Coates et al., 2018). For example, the identification of the narrowly distributed *N. vittata* [B] lineage implies that it may be of greater conservation concern than the current status of *N. vittata* would denote (Buckley et al., 2018), with its situation being more parallel to *N. pygmaea* considered as one of the most imperilled freshwater fishes in the country (Lintermans et al., 2020). Given that *N. vittata* [B] and *N. pygmaea* occupy narrow distributions under threat from salinisation and aridification (Allen et al., 2020), and that pygmy perches are highly vulnerable to human-induced habitat loss (Brauer & Beheregaray, 2020), *ex situ* conservation efforts such as translocation and the formation of insurance populations may be required in the immediate future (as done for other pygmy perches; e.g. Marshall et al., 2022).

A previous phylogenomic study of all pygmy perch species, including species delimitation using a multi-species coalescent approach, suggested that *N. vittata* was a complex of three cryptic species (Buckley et al., 2018). However, this was based on few populations within SWWA, all of which were included in this study. Two of these species correspond to the most divergent populations within *N. vittata* [A] (NviCBP and NviAR), suggesting that the higher number of species proposed in the previous study reflected a lack of intermediate sampling within the lineage. However, given that NviCBP was the most divergent population within the *N. vittata* [A] lineage despite being located in the centre of its geographic range, the possibility remains that it represents a fourth candidate species, as predicted by an allozyme dataset for over 40 *N. vittata* [A] sites (M. Adams, unpublished).

### Phylogeographic Structure Within the SWWA

Intraspecific phylogeographic structure was also detected in the most widespread putative species (*N. vittata* [A]), with three groups possibly representing evolutionarily-significant units (ESU) following Funk, McKay, Hohenlohe, & Allendorf (2012). This is based on their relatively ancient divergence (>1 Myr) and long-term demographic and spatial isolation. Within this delineation, the Margaret River population (NviCBP) represented a unique ESU: this is corroborated by other aquatic taxa in the region, with a unique ESU of a freshwater mussel (*Westralunio carteri;* Benson, Stewart, Close, & Lymbery, 2022) and two morphologically distinct species of freshwater crayfish (the hairy marron (*Cherax tenuimanus;* Vercoe, Lawrence, & de Graaf, 2009) and the Margaret River burrowing crayfish (*Engaewa pseudoreducta*; Allen, Beatty & Morgan, 2017)) endemic to Margaret River. The remaining two ESUs captured disjunct pairs of populations at opposing extremes of the species distribution (in the north and the southeast).

### Implications for Conservation of a Biodiversity Hotspot

Freshwater biodiversity is considered one of the most threatened groups globally (Collen et al., 2014; Lintermans et al., 2020). Within the SWWA, freshwater species are currently threatened by several convergent issues, including ongoing aridification since the mid-last century (Smith & Power, 2014), habitat clearing primarily for agricultural development (Andrich & Imberger, 2013), secondary salinization of rivers (Morgan, Thorburn & Gill, 2003; Allen et al., 2020) and invasive species (Morgan, Gill, Maddern & Beatty, 2004; Beatty & Morgan, 2013). These threats will likely be exacerbated by anthropogenic climate change and their own interactive effects (Beatty, Morgan, & Lymbery, 2014; Stewart, Ford, & Benson, 2022): projections of climate change alterations alone predict a 40–50% decrease in plant ranges and 10–44% loss of endemic plant diversity (294–1,293 species) in the SWWA (Habel et al., 2019). Additionally, spatial heterogeneity in the effects of climate change may affect some lineages more than others (Hansen et al., 2019; Stewart et al., 2022), such as due to a northern contraction of suitable mesic habitat (Klausmeyer & Shaw, 2009) or loss of permanent refuge pools (Allen et al., 2020). Relatedly, a reduction in fitness associated with heterogeneous climate change has already been observed in some SWWA plant species (Brouwers et al., 2012; Dalmaris et al., 2015; Monks et al., 2019). Our projections of species distributions under climate change echo these observations, suggesting a northern and inland contraction of suitable habitat for *N. vittata.* Although this would likely entail a range reduction of the more widespread putative species (*N. vittata* [A]), a LDM of *N. vittata* [B] similarly predicted substantial (potentially total) range contraction as well. Conservation programs must therefore consider both the spatial heterogeneity of these impacts, and how they may interact in increasing extinction risk, especially for freshwater fauna.

Disruption of climatic regimes, in conjunction with broadly increased temperatures (Jeremias et al., 2018), aridity (Davis, Pavlova, Thompson, & Sunnucks, 2013) and sea levels (Rotzoll & Fletcher, 2012) is already impacting species persistence globally. These effects will be exacerbated within regions of low historical climate variability due to both low species resilience and high species diversity, with compounding effects for coastal temperate ecosystems (Buckley et al., 2021).

Additionally, the high biodiversity of historically stable regions may mean they become hotspots of extinction as swathes of taxa struggle to respond to the selective challenges of anthropogenic climate change (Waldvogel et al., 2020). Given that biodiversity hotspots may also have an excess of undiscovered and cryptic biodiversity (Joppa et al., 2011), current estimates of extinction rates may be underestimated (Le Roux et al., 2019). Two of the three putative species delineated here have small ranges, a pattern shared by other narrow-range endemics in biodiversity hotspots (Goldberg, Roy, Lande, & Jablonski, 2005). Together, these components highlight why biodiversity hotspots are regions of high conservation concern and should be priorities in conservation management to maximise conservation of at-risk species and ecological communities.

Long-term climatic stability has driven the accumulation of divergent and isolated lineages of freshwater fishes in SWWA, providing a mechanism for its status as a biodiversity hotspot. This included multiple diversifications within the *N. vittata* complex. Given that regional climates are being disrupted globally by climate change, and that species that evolved in historically stable regions may lack the adaptive mechanisms to respond, biodiversity hotspots like SWWA may be exceptionally threatened. These findings only further solidify the conservation need for these regions, with preservation of threatened habitat such as rivers and refugial pools crucial for the persistence of a suite of endemic species. For those species most at risk, adaptive conservation management strategies such as the formation of insurance populations and captive breeding programs may become necessary to protect lineages with lower adaptive resilience to rapidly changing environments.

## ACKNOWLEDGMENTS

This research was supported by Australian Research Council grants (FT130101068 and DP190102533) to LBB and an Australian Government Research Training Program Scholarship to SJB. This work received logistic support from Flinders University, University of Canberra, Murdoch University and the South Australian Museum. We additionally thank Michael S. Y. Lee for assistance with species delimitation analyses. The authors acknowledge the Noongar people who are the Traditional Custodians of the land on which this research took place.

## AUTHOR CONTRIBUTIONS

SJB contributed to all sections of data generation and analysis, as well as drafting the manuscript. LBB designed and supervised the study, and helped with manuscript drafting. PU, MH, MA, SB and DM all contributed with samples and field expertise. CB assisted with population genomic analyses and interpretation. All authors contributed to the interpretation of results and critically revised the manuscript. All authors read and approved the final manuscript.

## DATA ARCHIVING

The data used in this study is available for peer-review at https://datadryad.org/stash/share/WHhKm5YTHxMQKoiiBfEq4cNvKtAWr6iJ_FhUECuVnKI.

## RESEARCH ETHICS STATEMENT

Collection of tissue samples were carried out in accordance with relevant guidelines and regulations of Australia and under Flinders University Animal Welfare Committee approval E313.

## References

Adams M, Raadik TA, Burridge CP, Georges A (2014) Global biodiversity assessment and hyper-cryptic species complexes: more than one species of elephant in the room? Syst Biol 63:518–533

Addo-Bediako A, Chown SL, Gaston KJ (2000) Thermal tolerance, climatic variability and latitude. Proc R Soc B Biol Sci 267:739–745

Ali R, McFarlane D, Varma S, Dawes W, Emelyanova I, Hodgson G et al. (2012) Potential climate change impacts on groundwater resources of south-western Australia. J Hydrol 475:456–472

Allen MG, Beatty SJ, Morgan DL (2017) Aquatic fauna refuges in Margaret River and the Cape to Cape region of Australia’s Mediterranean-climatic Southwestern Province. FiSHMED: Fishes in Mediterranean Environments, 2017.002, 1–26. doi:10.29094/FiSHMED.2017.002

Allen MG, Morgan DL, Close PG, Beatty SJ (2020) Too little but not too late? Biology of a recently discovered and imperilled freshwater fish in a drying temperate region and comparison with sympatric fishes. Aquat Conserv: Mar Freshw Ecosyst 30:1412– 1423

Andrich MA, Imberger J (2013) The effect of land clearing on rainfall and fresh water resources in Western Australia: a multi-functional sustainability analysis. Int J Sustain Dev World Ecol 20: 549–563

Baele G, Lemey P, Bedford T, Rambaut A, Suchard MA., Alekseyenko, AV (2012) Improving the accuracy of demographic and molecular clock model comparison while accommodating phylogenetic uncertainty. Mol Biol Evol 29:2157–2167

Beatty SJ, Morgan DL (2013) Introduced freshwater fishes in a global endemic hotspot and implications of habitat and climatic change. Bioinvasions Rec 2:1–9

Beatty SJ, Morgan, DL, Lymbery AJ (2014) Implications of climate change for potamodromous fishes. Glob Change Biol 20:1794–1807

Beatty SJ, Morgan DL, Rashnavadi M, Lymbery AJ (2011) Salinity tolerances of endemic freshwater fishes of south-western Australia: implications for conservation in a biodiversity hotspot. Mar Freshw Res 62:91–100

Benson J, Stewart B, Close P, Lymbery A (2022) Evidence for multiple refugia and hotspots of genetic diversity for *Westralunio carteri*, a threatened freshwater mussel in south-western Australia. Aquat Conserv: Mar Freshw Ecosyst 32:559–575

Bi D, Dix M, Marsland S, O’Farrell S, Sullivan A, Bodman R et al. (2020) Configuration and spin-up of ACCESS-CM2, the new generation Australian Community Climate and Earth System Simulator Coupled Model. J South Hemisphere Earth Syst Sci 70:225– 251

Bouckaert R, Vaughan T G, Barido-Sottani J, Duchêne S, Fourment M, Gavryushkina A et al. (2019) BEAST 2.5: An advanced software platform for Bayesian evolutionary analysis. PLoS Comput Biol 15:e1006650

Brauer CJ, Beheregaray LB (2020) Recent and rapid anthropogenic habitat fragmentation increases extinction risk for freshwater biodiversity. Evol Appl 13:2857–2869

Brauer CJ, Hammer MP, Beheregaray LB (2016) Riverscape genomics of a threatened fish across a hydroclimatically heterogeneous river basin. Mol Ecol 25:5093–5113

Brouwers NC, Mercer J, Lyons T, Poot P, Veneklaas E, Hardy G (2012) Climate and landscape drivers of tree decline in a Mediterranean ecoregion. Ecol Evol 3:67–79

Brown JL, Bennett JR, French CM (2017) SDMtoolbox 2.0: the next generation Python-based GIS toolkit for landscape genetic, biogeographic and species distribution model analyses. PeerJ 5:e4095

Brown JL, Hill DJ, Dolan AM, Carnaval AC, Haywood AM (2018) PaleoClim, high spatial resolution paleoclimate surfaces for global land areas. Sci Data 5:180254

Bryant D, Bouckaert R, Felsenstein J, Rosenberg NA, RoyChoudhury A (2012) Inferring species trees directly from biallelic genetic markers: Bypassing gene trees in a full coalescent analysis. Mol Biol Evol 29:1917–1932

Buckley SJ, Brauer CJ, Unmack PJ, Hammer MP, Beheregaray LB (2021) The roles of aridification and sea level changes in the diversification and persistence of freshwater fish lineages. Mol Ecol 30:4866–4883

Buckley SJ, Domingos FMCB, Attard C, Brauer CJ, Sandoval-Castillo J, Lodge R et al. (2018) Phylogenomic history of enigmatic pygmy perches: implications for biogeography, taxonomy and conservation. Royal Soc Open Sci, 5:172125

Byrne M, Steane DA, Joseph L, Yeates DK, Jordan GJ, Crayn D et al. (2011) Decline of a biome: evolution, contraction, fragmentation, extinction and invasion of the Australian mesic zone biota. J Biogeogr 38:1635–1656

Carnaval AC, Hickerson MJ, Haddad CB, Rodrigues MT, Moritz C (2009) Stability predicts genetic diversity in the Brazilian Atlantic forest hotspot. Sci 323:785–789

Catchen J, Hohenlohe PA, Bassham S, Amores A, Cresko WA (2013) Stacks: an analysis tool set for population genomics. Mol Ecol 22:3124–3140

Chambers EA, Hillis DM (2020) The multispecies coalescent over-splits species in the case of geographically widespread taxa. Syst Biol 69:184–193

Coates DJ, Byrne M, Moritz C (2018) Genetic diversity and conservation units: Dealing with the species-population continuum in the age of genomics. Front Ecol Evol 6:165

Collen B, Whitton F, Dyer EE, Baillie JE, Cumberlidge N, Darwall WR et al. (2014) Global patterns of freshwater species diversity, threat and endemism. Glob Ecol Biogeogr 23:40–51.

Cowling RM, Lombard AT (2002) Heterogeneity, speciation/extinction history and climate: explaining regional plant diversity patterns in the Cape Floristic Region. Divers Distrib 8:163–179

Crisp MD, Cook LG (2007) A congruent molecular signature of vicariance across multiple plant lineages. Mol Phylogenet Evol 43:1106–1117

Dalmaris E, Ramalho CE, Poot P, Veneklaas EJ, Byrne M (2015) A climate change context for the decline of a foundation tree species in south-western Australia: insights from phylogeography and species distribution modelling. Ann Bot 116:941–952

Davis J, Pavlova A, Thompson R, Sunnucks P (2013) Evolutionary refugia and ecological refuges: key concepts for conserving Australian arid zone freshwater biodiversity under climate change. Glob Change Biol 19:1970–1984

Derkarabetian S, Castillo S, Koo PK, Ovchinnikov S, Hedin M (2019) A demonstration of unsupervised machine learning in species delimitation. Mol Phylogenet Evol 139:106562

Drummond AJ, Rambaut A (2019) TreeAnnotator (Version 2.6). http://beast.bio.ed.ac.uk.

Eaton DA (2014). PyRAD: assembly of de novo RADseq loci for phylogenetic analyses. Bioinformatics 30:1844–1849

Elith J, Phillips SJ, Hastie T, Dudík M, Chee YE, Yates CJ (2011) A statistical explanation of MaxEnt for ecologists. Divers Distrib 17:43–57

Fick SE, Hijmans RJ (2017) WorldClim 2: new 1-km spatial resolution climate surfaces for global land areas. Int J Climatol 37:4302 – 4315

Funk WC, McKay JK, Hohenlohe PA, Allendorf FW (2012) Harnessing genomics for delineating conservation units. Trends Ecol Evol 27:489–496

Funnekotter AV, Millar M, Krauss SL, Nevill PG (2019) Phylogeographic analyses of *Acacia karina* (Fabaceae) support long term persistence of populations both on and off banded iron formations. Aust J Bot 67:194

Gioia P, Hopper SD (2017) A new phytogeographic map for the Southwest Australian Floristic Region after an exceptional decade of collection and discovery. Bot J Linn Soc 184:1–15

Goldberg EE, Roy K, Lande R, Jablonski D (2005) Diversity, endemism, and age distributions in macroevolutionary sources and sinks. Am Nat 165:623–633

Gruber B, Unmack PJ, Berry OF, Georges A (2018) dartr: An r package to facilitate analysis of SNP data generated from reduced representation genome sequencing. Mol Ecol Resour 18:691–699

Grummer JA, Bryson Jr. RW, Reeder TW (2013) Species delimitation using Bayes factors: Simulations and application to the *Sceloporus scalaris* species group (Squamata: Phrynosomatidae). Syst Biol 63:119–133

Habel JC, Rasche L, Schneider UA, Engler JO, Schmid E, Rödder D et al. (2019) Final countdown for biodiversity hotspots. Conserv Lett 12

Hallett CS, Hobday AJ, Tweedley JR, Thompson PA, McMahon K, Valesini FJ (2018). Observed and predicted impacts of climate change on the estuaries of south-western Australia, a Mediterranean climate region. Reg Environ Change 18:1357–1373

Hansen BB, Pedersen ÅØ, Peeters B, Le Moullec M, Albon SD, Herfindal I et al. (2019) Spatial heterogeneity in climate change effects decouples the long-term dynamics of wild reindeer populations in the high Arctic. Glob Change Biol 25:3656–3668

Hao T, Elith J, Guillera-Arroita G, Lahoz-Monfort JJ (2019) A review of evidence about use and performance of species distribution modelling ensembles like BIOMOD. Divers Distrib 25:839–852

Harrison S, Noss R (2017) Endemism hotspots are linked to stable climatic refugia. Ann Bot 119:207–214

Hopper SD, Gioia P (2004) The Southwest Australian Floristic Region: Evolution and conservation of a global hot spot of biodiversity. Annu Rev Ecol Evol Syst 35:623– 650

Jeremias G, Barbosa J, Marques SM, Asselman J, Gonçalves FJM, Pereira JL (2018) Synthesizing the role of epigenetics in the response and adaptation of species to climate change in freshwater ecosystems. Mol Ecol 27:2790–2806

Joppa LN, Roberts DL, Myers N, Pimm SL (2011) Biodiversity hotspots house most undiscovered plant species. Proc Natl Acad Sci U.S.A. 108:13171–13176

Klausmeyer KR, Shaw MR (2009) Climate change, habitat loss, protected areas and the climate adaptation potential of species in mediterranean ecosystems worldwide. PLoS One 4:e6392

Le Roux JJ, Hui C, Castillo ML, Iriondo JM, Keet JH, Khapugin AA et al. (2019) Recent anthropogenic plant extinctions differ in biodiversity hotspots and coldspots. Curr Biol 29:2912–2918

Leaché AD, Fujita MK, Minin VN, Bouckaert RR (2014) Species delimitation using genome-wide SNP data. Syst Biol 63:534–542

Lintermans M, Geyle HM, Beatty S, Brown C, Ebner BC, Freeman R et al. (2020) Big trouble for little fish: identifying Australian freshwater fishes in imminent risk of extinction. Pac Conserv Biol 26:365–377

Liu L, Xi Z, Davis CC (2015) Coalescent methods are robust to the simultaneous effects of long branches and incomplete lineage sorting. Mol Biol Evol 32:791–805

Malinsky M, Matschiner M, Svardal H (2021) Dsuite - fast D-statistics and related admixture evidence from VCF files. Mol Ecol Resour 21:584–595

Marmion M, Parviainen M, Luoto M, Heikkinen RK, Thuiller W (2009) Evaluation of consensus methods in predictive species distribution modelling. Divers Distrib 15:59– 69

Marshall IR, Brauer CJ, Wedderburn SD, Whiterod NS, Hammer MP, Barnes TC, Attard CRM, Möller LM, Beheregaray LB (2022) Longitudinal monitoring of neutral and adaptive genomic diversity in a reintroduction. Conserv Biol 36:e13889

Martin SH, Davey JW, Jiggins CD (2014) Evaluating the use of ABBA–BABA statistics to locate introgressed loci. Mol Biol Evol 32:244–257

Minh BQ, Hahn MW, Lanfear R (2020a) New methods to calculate concordance factors for phylogenomic datasets. Mol Biol Evol 37:2727–2733

Minh BQ, Schmidt HA, Chernomor O, Schrempf D, Woodhams MD, von Haeseler A et al. (2020b) IQ-TREE 2: New models and efficient methods for phylogenetic inference in the genomic era. Mol Biol Evol 37:1530–1534

Moise A, Bhend J, Watterson I, Wilson L (2015) Evaluation of climate models. In: Whetton P (ed) Climate Change in Australia Information for Australia’s Natural Resource Management Regions: Technical Report. CSIRO and Bureau of Metereology, Australia, pp 53–78

Monks L, Barrett S, Beecham B, Byrne M, Chant A, Coates D et al. (2019) Recovery of threatened plant species and their habitats in the biodiversity hotspot of the Southwest Australian Floristic Region. Plant Divers 41:59–74

Morgan DL, Beatty SJ, Adams M (2013) *Nannoperca pygmaea*, a new species of pygmy perch (Teleostei: Percichthyidae) from Western Australia. Zootaxa 3637:401

Morgan DL, Gill HS, Maddern MG, Beatty SJ (2004) Distribution and impacts of introduced freshwater fishes in Western Australia. N. Z. J Mar Freshwater Res 38:511–523.

Morgan DL, Gill HS, Potter IC (1998) Distribution, identification and biology of freshwater fishes in south-western Australia. Rec West Aust Mus, Suppl 56:1–97

Morgan DL, Thorburn DC, Gill HS (2003) Salinization of south-western Western Australian rivers and the implications for the inland fish fauna – the Blackwood River, a case study. Pac Conserv Biol 9:161–171

Myers N, Mittermeier RA, Mittermeier CG, da Fonseca GAB, Kent J (2000) Biodiversity hotspots for conservation priorities. Nature 403:853–858

Nevill PG, Bradbury D, Williams A, Tomlinson S, Krauss SL (2014) Genetic and palaeo-climatic evidence for widespread persistence of the coastal tree species *Eucalyptus gomphocephala* (Myrtaceae) during the Last Glacial Maximum. Ann Bot 113:55–67

Nistelberger H, Gibson N, Macdonald B, Tapper SL, Byrne M (2014) Phylogeographic evidence for two mesic refugia in a biodiversity hotspot. Heredity 113:454–463

Ogsten G, Beatty SJ, Morgan DL, Pusey B, Lymbery AJ (2016) Living on burrowed time: Aestivating fishes in south-western Australia face extinction due to climate change. Biol Conserv 195:235–244

Patterson N, Moorjani P, Luo Y, Mallick S, Rohland N, Zhan Y et al. (2012) Ancient admixture in human history. Genetics 192:1065–1093

Pickrell JK, Pritchard JK (2012) Inference of population splits and mixtures from genome-wide allele frequency data. PLoS Genet 8:e1002967

Pittock J, Hansen LJ, Abell R (2008) Running dry: Freshwater biodiversity, protected areas and climate change. Biodivers 9:30–38

Raftery, A. E., Newton, M. A., Satagopan, J. M., & Krivitsky, P. N. (2006). Estimating the integrated likelihood via posterior simulation using the harmonic mean identity. Bayesian Statistics.

Rambaut A, Drummond AJ (2009). Tracer v1.5. http://beast.bio.ed.ac.uk.

Rix MG, Edwards DL, Byrne M, Harvey MS, Joseph L, Roberts JD (2015) Biogeography and speciation of terrestrial fauna in the south-western Australian biodiversity hotspot. Biol Rev 90:762–793

Rix MG, Harvey MS (2012) Phylogeny and historical biogeography of ancient assassin spiders (Araneae: Archaeidae) in the Australian mesic zone: evidence for Miocene speciation within Tertiary refugia. Mol Phylogenet Evol 62:375–396

Rotzoll K, Fletcher CH (2012) Assessment of groundwater inundation as a consequence of sea-level rise. Nat Clim Change 3:477

Sandel B, Arge L, Dalsgaard B, Davies RG, Gaston KJ, Sutherland WJ, Svenning J-C (2011).The influence of Late Quaternary climate-change velocity on species endemism. Science 334:660–664

Sandoval-Castillo J, Gates K, Brauer CJ, Smith S, Bernatchez L, Beheregaray LB (2020) Adaptation of plasticity to projected maximum temperatures and across climatically defined bioregions. Proc Natl Sci Acad U. S. A. 117:17112–17121

Smith I, Power S (2014) Past and future changes to inflows into Perth (Western Australia) dams. J Hydrol: Reg Stud 2:84–96

Spooner MI, De Deckker P, Barrows TT, Fifield LK (2011) The behaviour of the Leeuwin Current offshore NW Australia during the last five glacial–interglacial cycles. Global Planet Change 75:119–132

Stamatakis A (2014). RAxML version 8: a tool for phylogenetic analysis and post-analysis of large phylogenies. Bioinf 30:1312–1313

Stange M, Sanchez-Villagra MR, Salzburger W, Matschiner M (2018) Bayesian divergence-time estimation with genome-wide single-nucleotide polymorphism data of sea catfishes (Ariidae) supports Miocene closure of the Panamanian Isthmus. Syst Biol 67:681–699

Stanton DWG, Frandsen P, Waples RK, Heller R, Russo I-RM, Orozco-terWengel PA et al. (2019) More grist for the mill? Species delimitation in the genomic era and its implications for conservation. Conserv Genetc 20:101–113

Stewart BA, Ford BJ, Benson JA (2022) Using species distribution modelling to identify ‘coldspots’ for conservation of freshwater fishes under a changing climate. Aquat Conserv Mar Freshwater Ecosyst 32:576–590

Stewart BA, Ford BJ, Van Helden BE, Roberts JD, Close PG, Speldewinde PC (2018) Incorporating climate change into recovery planning for threatened vertebrate species in southwestern Australia. Biodivers Conserv 27:147 – 165

Struck TH, Feder JL, Bendiksby M, Birkeland S, Cerca J, Gusarov VI et al. (2018) Finding evolutionary processes hidden in cryptic species. Trends Ecol Evol 33:153–163

Sundaram M, Donoghue MJ, Farjon A, Filer D, Mathews S, Jetz W, Leslie AB (2019) Accumulation over evolutionary time as a major cause of biodiversity hotspots in conifers. Proc R Soc B Biol Sci 286:20191887

Sunnucks P, Hales DF (1996) Numerous transposed sequences of mitochondrial cytochrome oxidase I-II in aphids of the genus *Sitobion* (Hemiptera: Aphididae). Mol Biol Evol 13:510–524

Swofford DL (2002). PAUP*. Phylogenetic Analysis Using Parsimony (*and Other Methods). Version 4.0b10 (Vol. Version 4.0). Sunderland, Massachusetts: Sinauer Associates.

Thuiller W, Lafourcade B, Engler R, Araújo MB (2009) BIOMOD – a platform for ensemble forecasting of species distributions. Ecography 32:369–373

Unmack PJ, Adams M, Hammer MP, Johnson JB, Gruber B, Gilles A et al. (2022) Plotting for change: an analytical framework to aid decisions on which lineages are candidate species in phylogenomic species discovery. Biol J Linn Soc 135:117–137

Unmack PJ, Hammer MP, Adams M, Dowling TE (2011) A phylogenetic analysis of pygmy perches (Teleostei: Percichthyidae) with an assessment of the major historical influences on aquatic biogeography in southern Australia. Syst Biol 60:797–812

Vercoe P, Lawrence C, de Graaf M (2009) Rapid replacement of the critically endangered hairy marron by the introduced smooth marron (Decapoda, Parastacidae) in the Margaret River (Western Australia). Crustaceana 82:1469–1476

Waldvogel AM, Feldmeyer B, Rolshausen G, Exposito-Alonso M, Rellstab C, Kofler R et al. (2020) Evolutionary genomics can improve prediction of species’ responses to climate change. Evol Lett 4:4–18

Waters JM, Burridge CP, Craw D (2020) River capture and freshwater biological evolution: A review of galaxiid fish vicariance. Diversity 12:216

Zhang C, Rabiee M, Sayyari E, Mirarab S (2018) ASTRAL-III: polynomial time species tree reconstruction from partially resolved gene trees. BMC Bioinf 19:153

