## Supplementary File 1 for "Long-term climatic stability drives accumulation and maintenance of divergent freshwater fish lineages in a temperate biodiversity hotspot"

**Table S1:** Locality data for *Nannoperca vittata* and *N. pygmaea* samples. Abbreviations described in the table refer to those used in further analyses and Figure 1. *Nannoperca obscura* samples were only included as an outgroup for phylogenetic analyses. *n* = number of individuals sequenced per population.

| Species | Putative species | Population | Abbreviation | Field code | Latitude | Longitude | *n* |
|---|---|---|---|---|---|---|---|
| *N. vittata* | *N. vittata* [A] | Arrowsmith R. | NviAR | FISHy6: DM146** | -29.626506 | 115.174854 | 4 |
| | | Preston R. | NviPre | E292: MA08-72NV | -33.3027 | 115.8176 | 4 |
| | | Angove R. | NviAng | FISHy4: EV-1:5 | -34.917 | 118.15 | 3 |
| | | Hay R. | NviHay | FISHy6: PU09-37NV | -34.834 | 117.406 | 4 |
| | | Canebrake Pool, Margaret R. | NviCBP | FISHy6: PU09-58NV | -33.880372 | 115.282701 | 2 |
| | *N. vittata* [B] | Doggerup Ck | NviDC | FISHy6: PU09-49NV | -34.741106 | 116.038006 | 4 |
| | | Chesapeake Brook | NviChes | FISHx2: V** | -34.833 | 116.333 | 4 |
| *N. pygmaea* | *N. pygmaea* | Mitchell R. | NpyMR | FISHy6: ESP0900* | -34.836211 | 117.412321 | 4 |
| | | Kent | NpyKent | FISHx2B: SB14 – SB23 | -34.6857 | 117.10272 | 4 |
| *N. obscura* | | Merri R., Grassmere | NobMRG | PU02-111YPP | -38.275 | 142.542 | 4 |
| Total | | | | 10 | | | 37 |

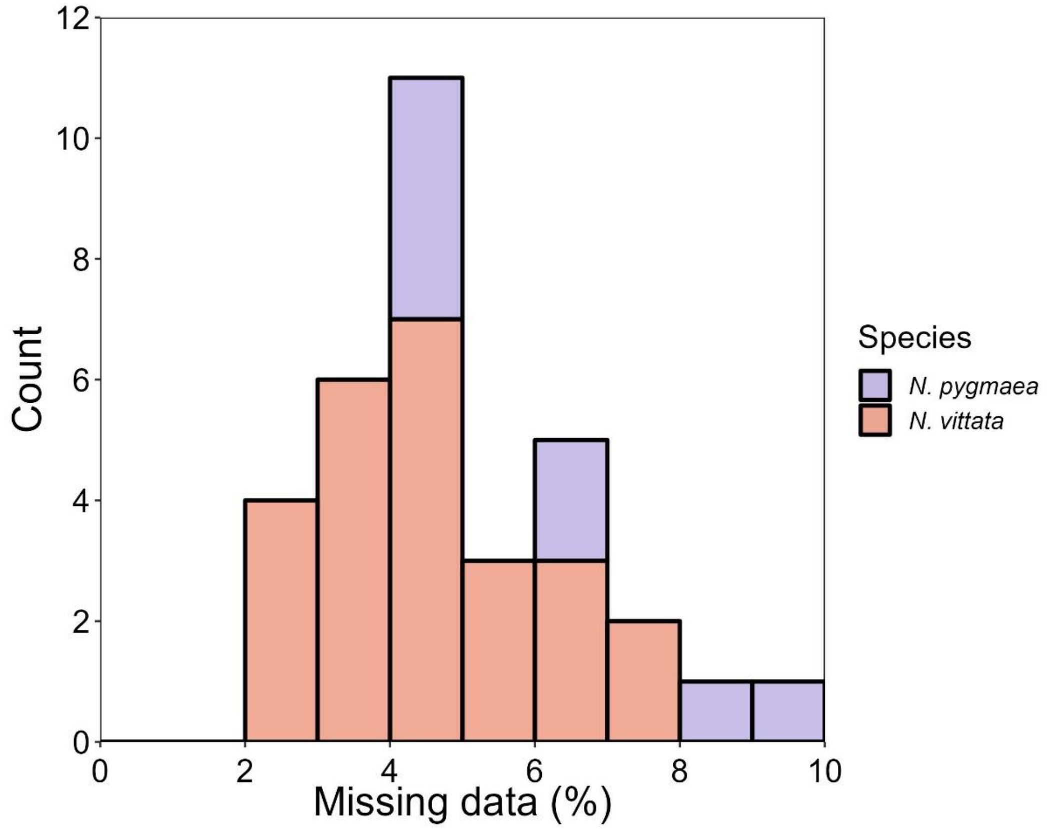


**Figure S1.** Stacked histogram of missing data per sample (%) for *Nannoperca vittata* and *N. pygmaea*.

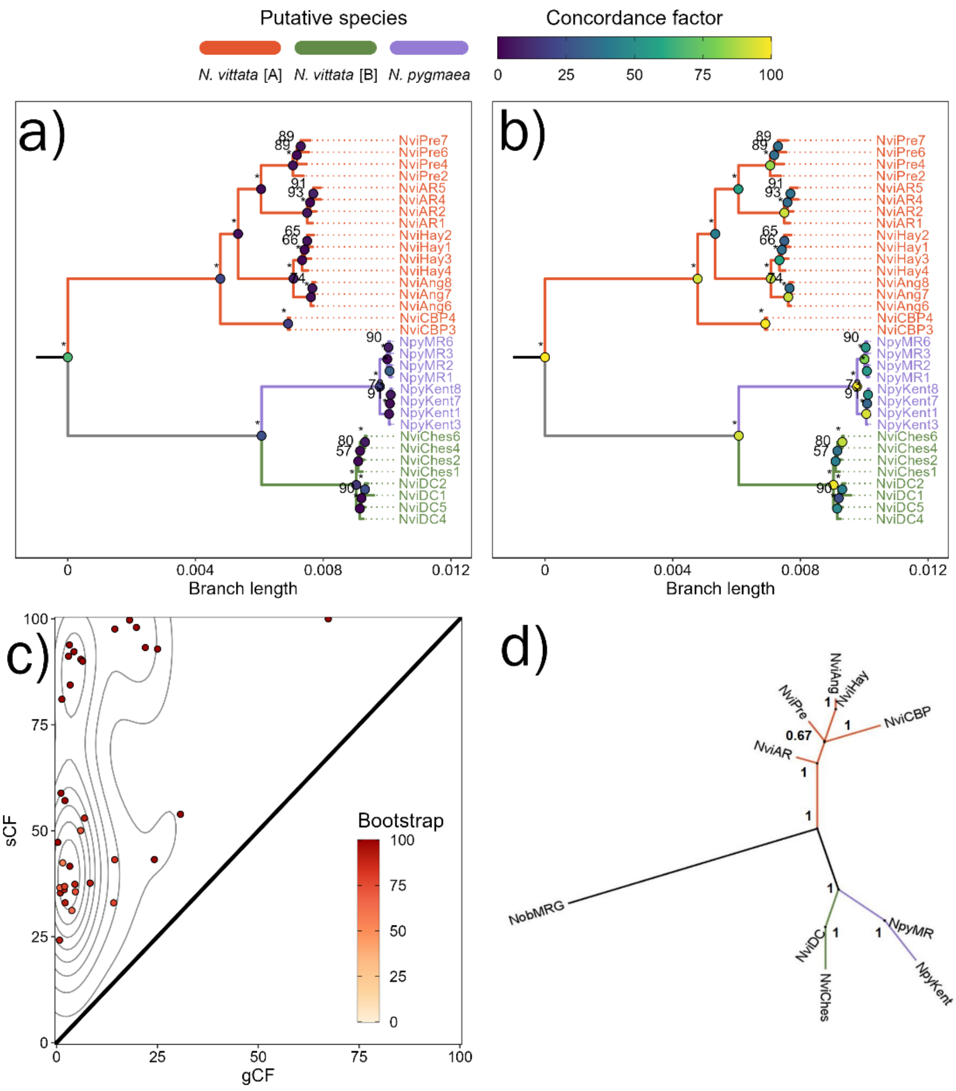


**Figure S2.** Summary of results from IQ-TREE2 and ASTRAL phylogenetic analyses. The phylogenetic trees in **a)** and **b)** represent the ML tree estimated by IQ-TREE2 with the alignment partitioned per ddRAD locus, with node labels showing bootstrap support. Node point colours denote gene (gCF) and site (sCF) concordance factors in **a)** and **b)**, respectively. Nodes with bootstrap support of 100 are indicated by asterisks. **c)** The relationship of partitioned bootstrap support, gCF and sCF. **d)** Species summary tree estimated by ASTRAL-III, with nodes labelled according to local posterior probability that branch is representative of the true species tree. For all phylogenetic plots, populations and branches are coloured according to putative species.

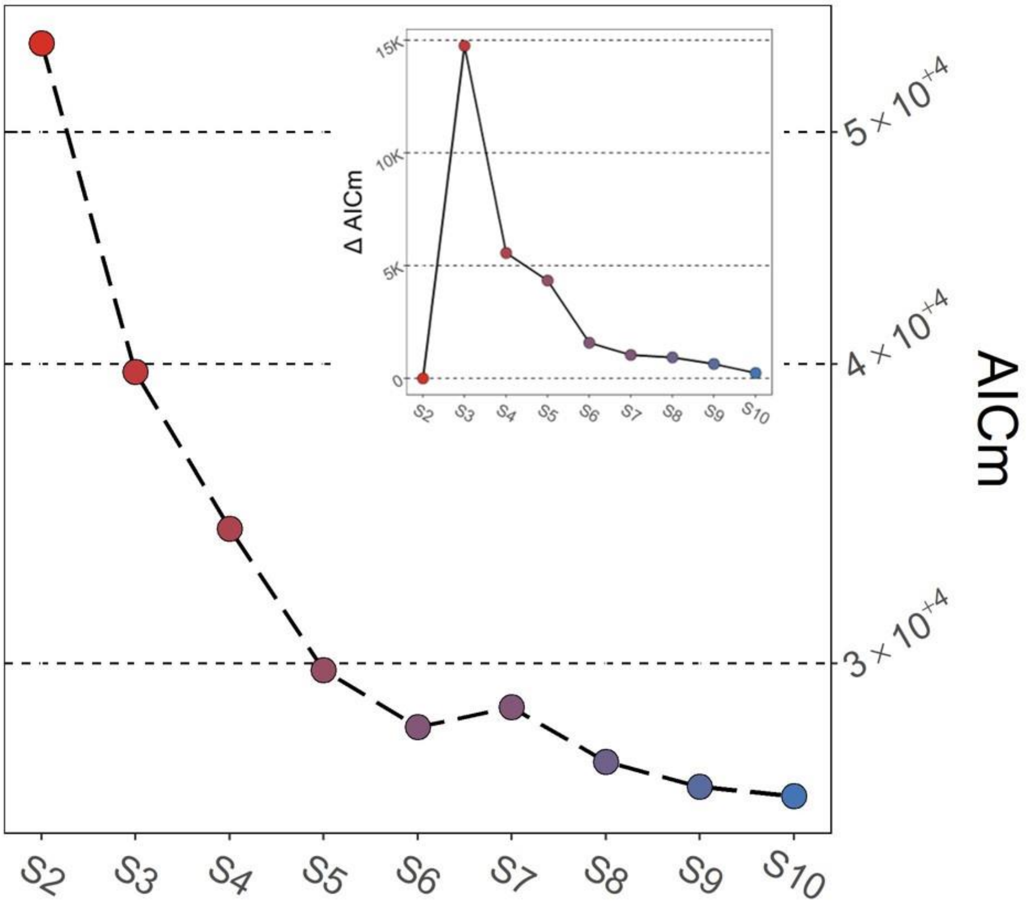


**Figure S3.** AICM values for species delimitation scenarios using SNAPP. Colours denote the number of species in the given model, ranging from two (red) to nine (blue). Inset depicts relative change in AICM with increasing number of species included.

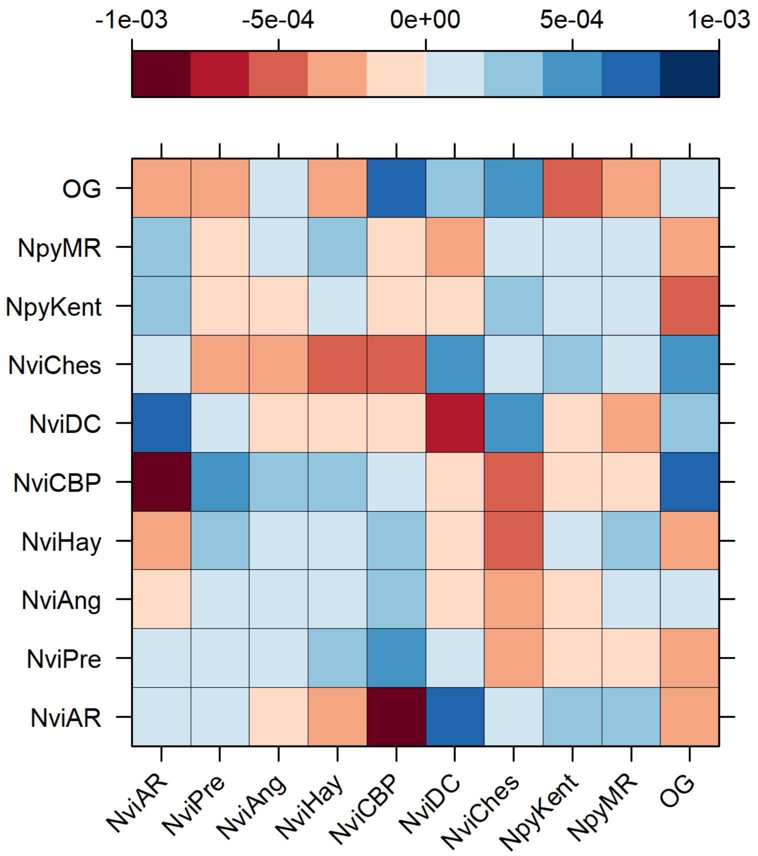


**Figure S4.** Covariance matrix of allele frequencies between populations without migration using TreeMix.

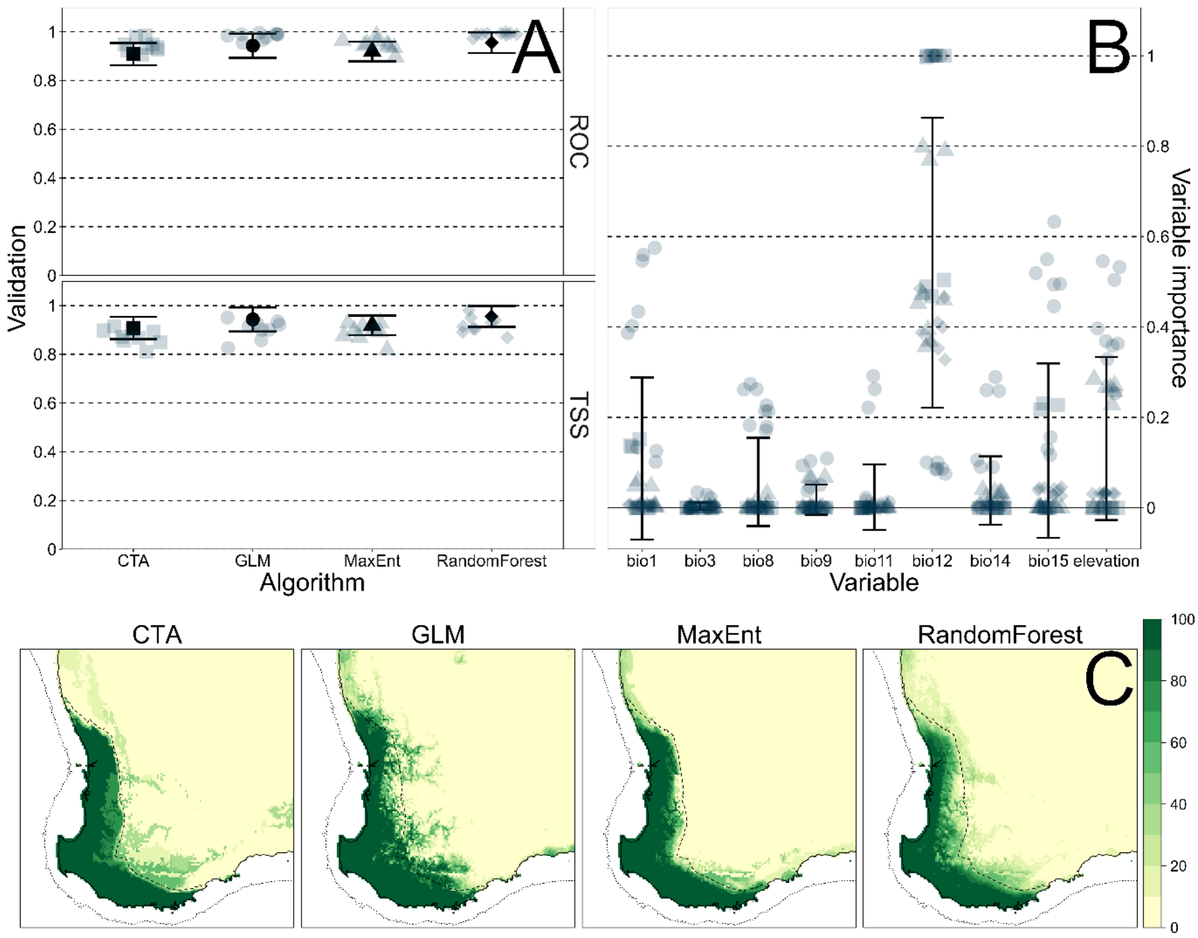


**Figure S5.** Evaluations of SDM accuracy. **a)** Model fit per algorithm (*n* = nine per algorithm) using the ROC and the TSS. **b)** Estimates of variable importance across all models (*n* = 36 total) for all variables used. Each point represents a single model (shapes indicate algorithm: see A) with error bars capturing the mean ± standard deviation across all models. **c)** Maps of ensemble distribution models built from averaging contemporary distributions for each method separately.

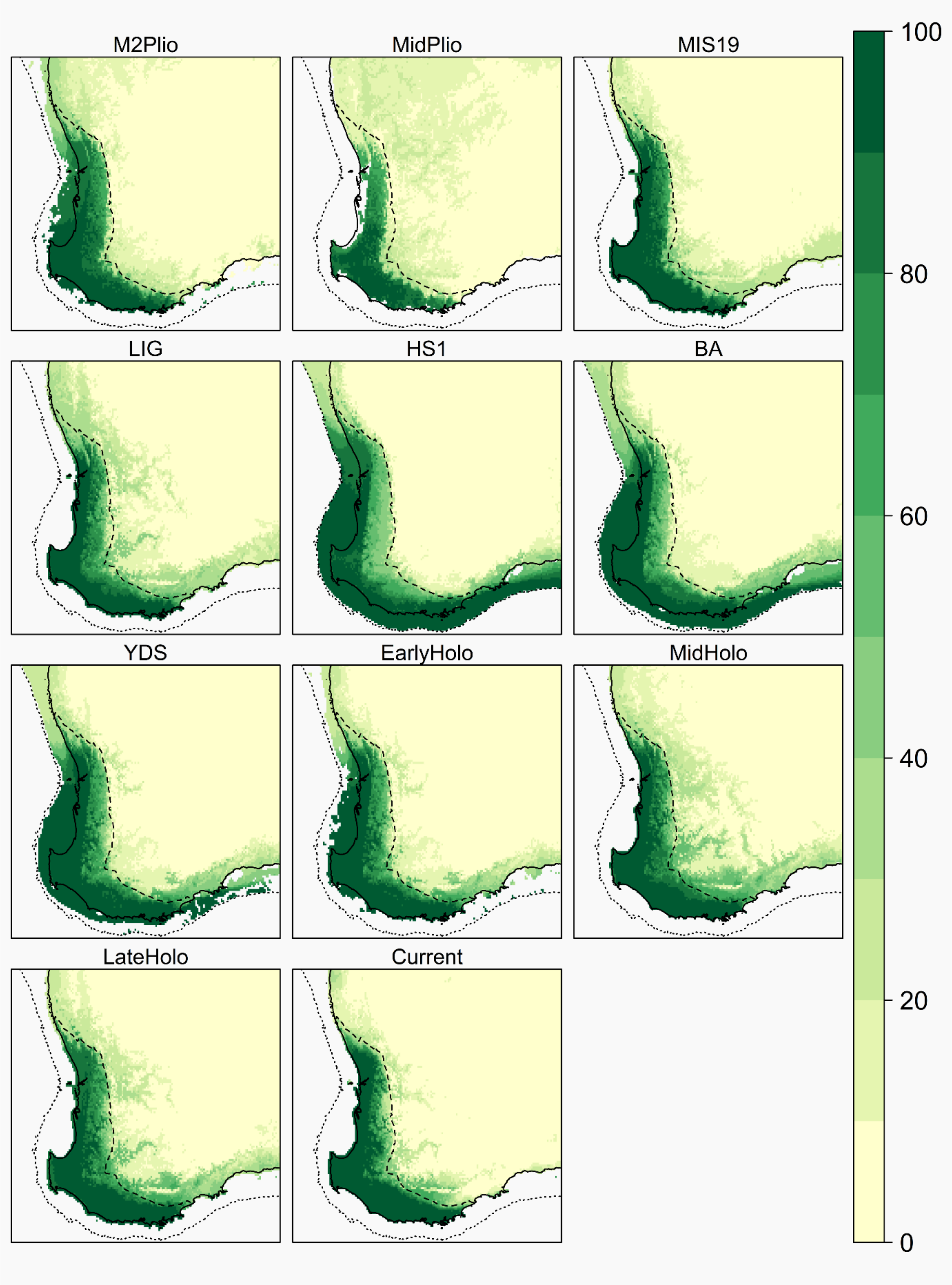


**Figure S6.** Historical projections of ensemble species distribution models. Maps are arranged in chronological order, from oldest (top left) to current (bottom right).

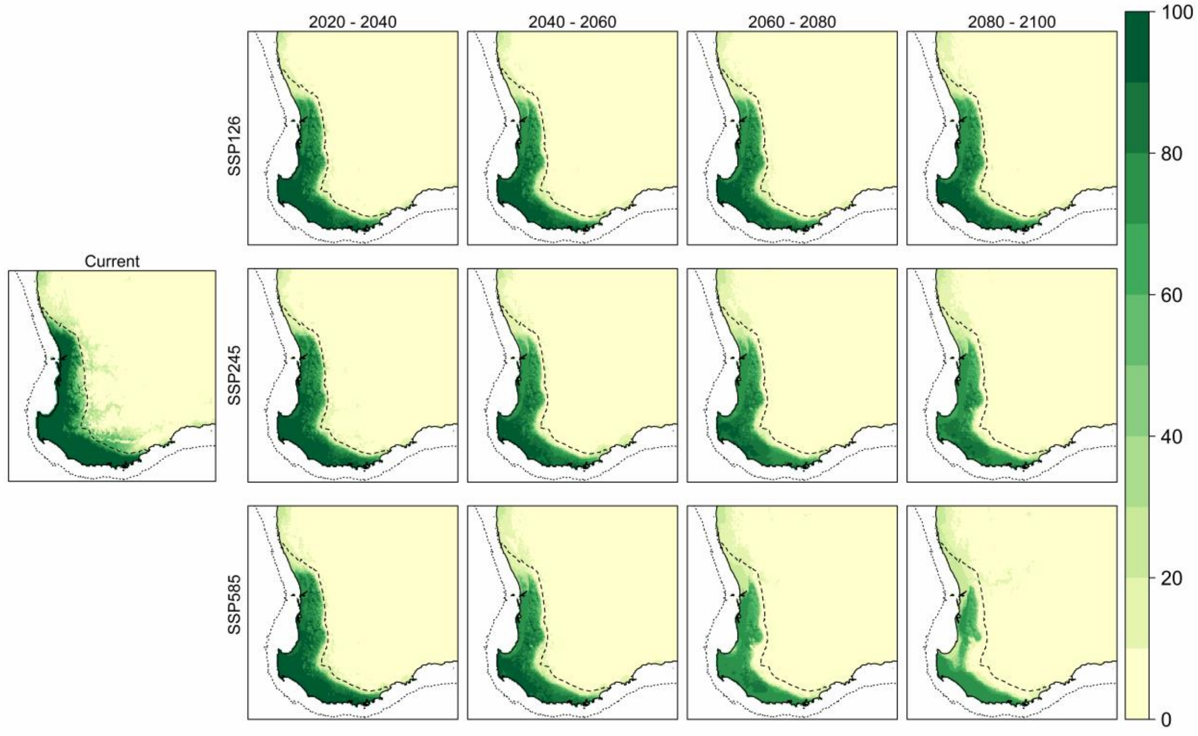


**Figure S7.** Future projections of ensemble species distribution models under climate change. Left: current ensemble distribution model. Map grid is separated by climate change scenarios (rows; from least severe at top to most severe at bottom) and time (from left to right).

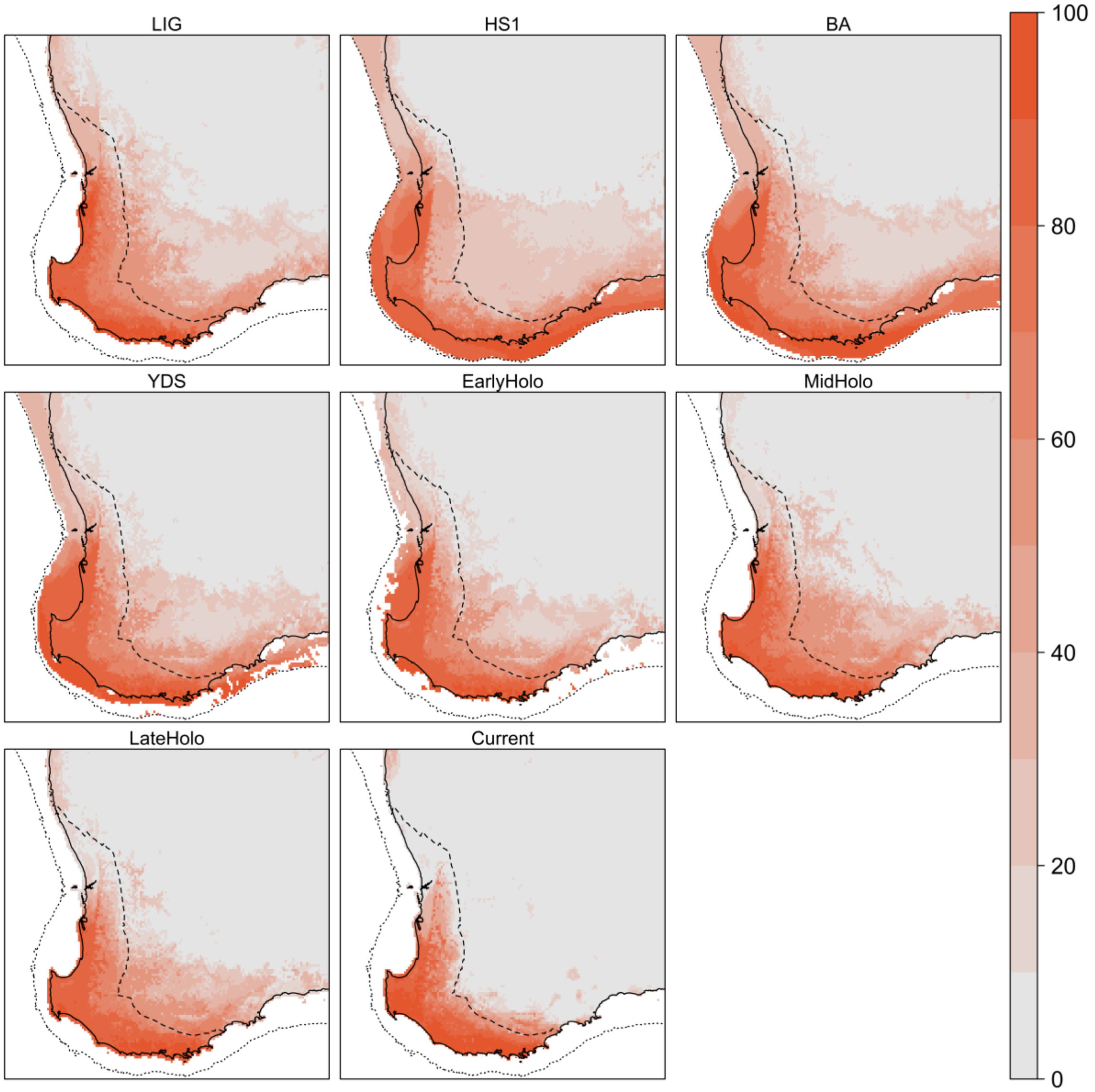


**Figure S8.** Historical projections of the lineage distribution model (LDM) for *N. vittata* [A]. Maps are arranged in chronological order, from oldest (top left) to current (bottom right).

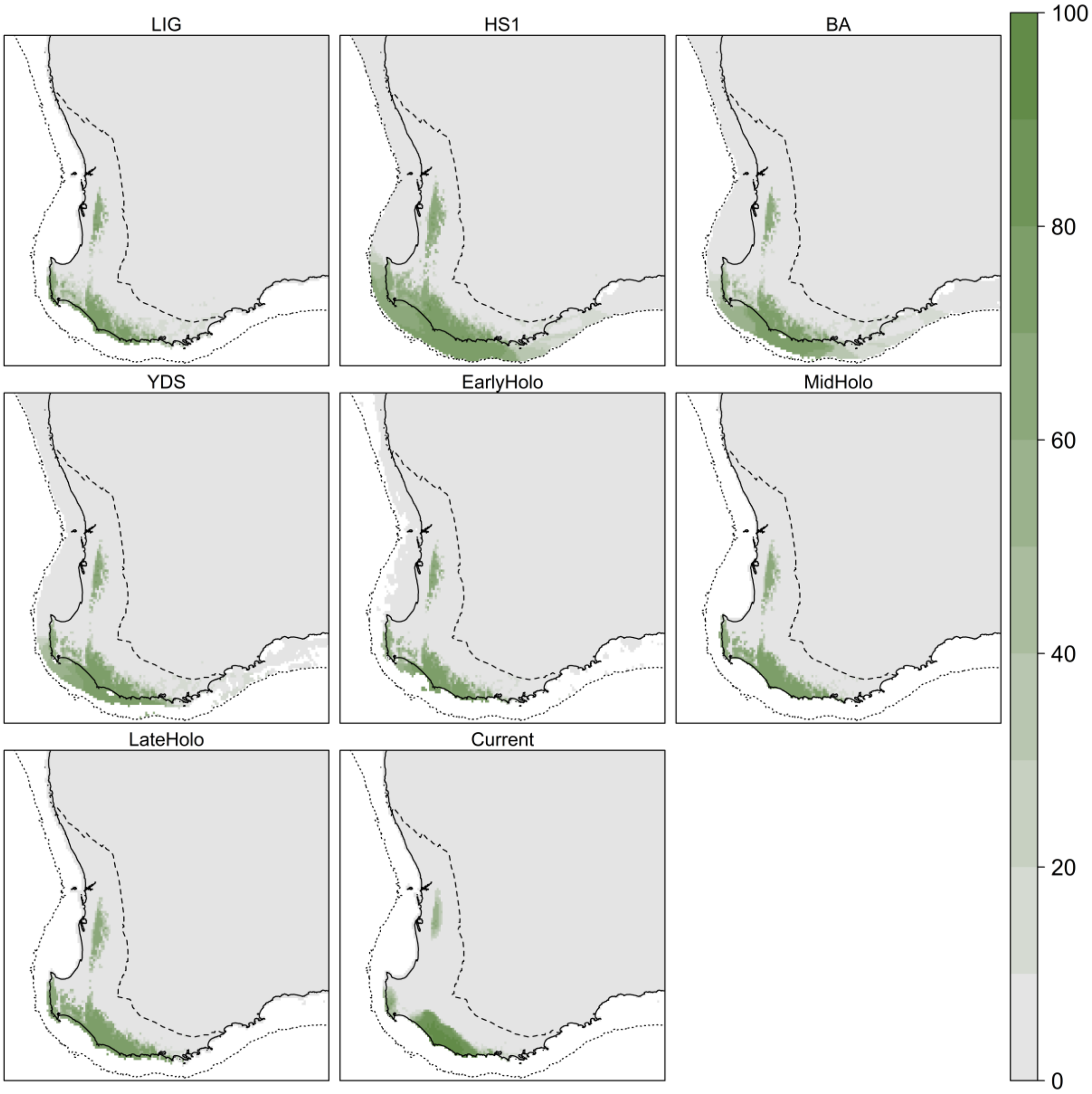


**Figure S9.** Historical projections of the lineage distribution model (LDM) for *N. vittata* [B]. Maps are arranged in chronological order, from oldest (top left) to current (bottom right).

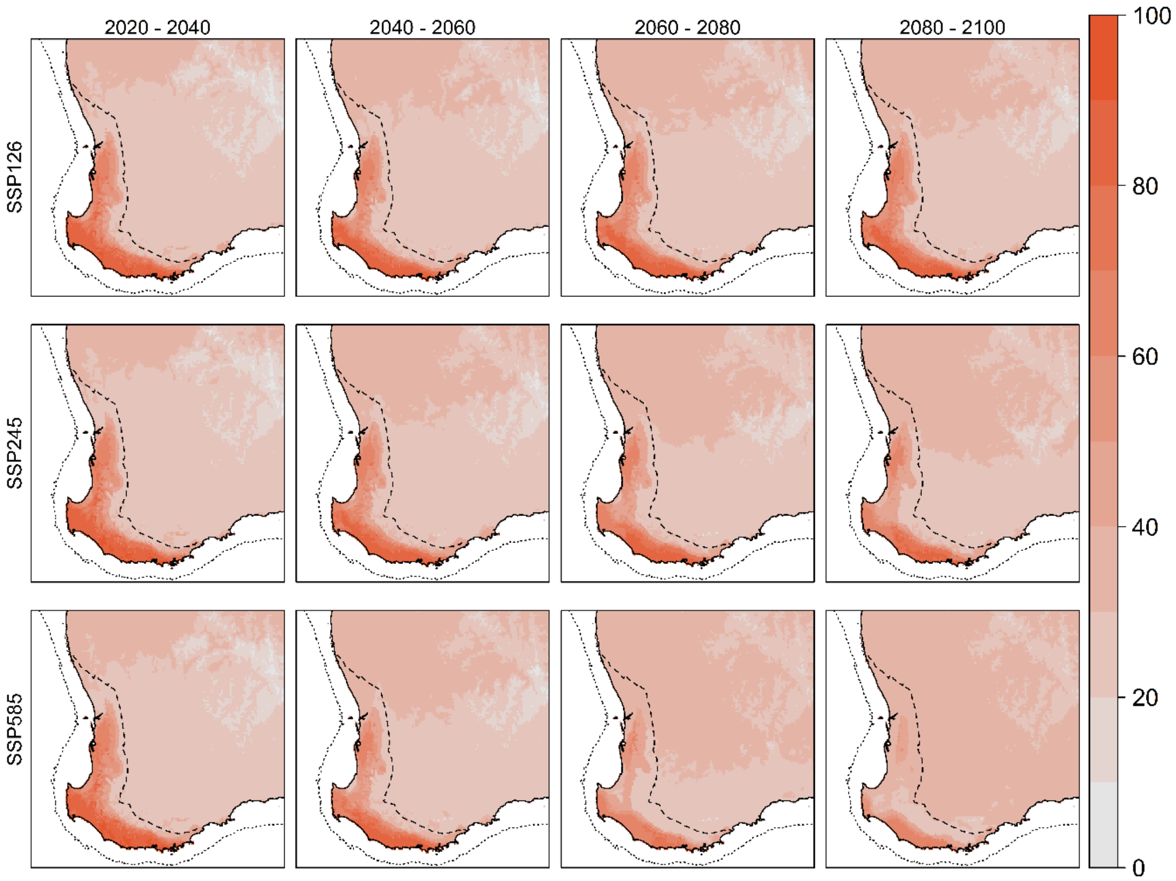


**Figure S10.** Future projections of the LDM for *N. vittata* [A] under climate change. Map grid is separated by climate change scenarios (rows; from least severe at top to most severe at bottom) and time (from left to right).

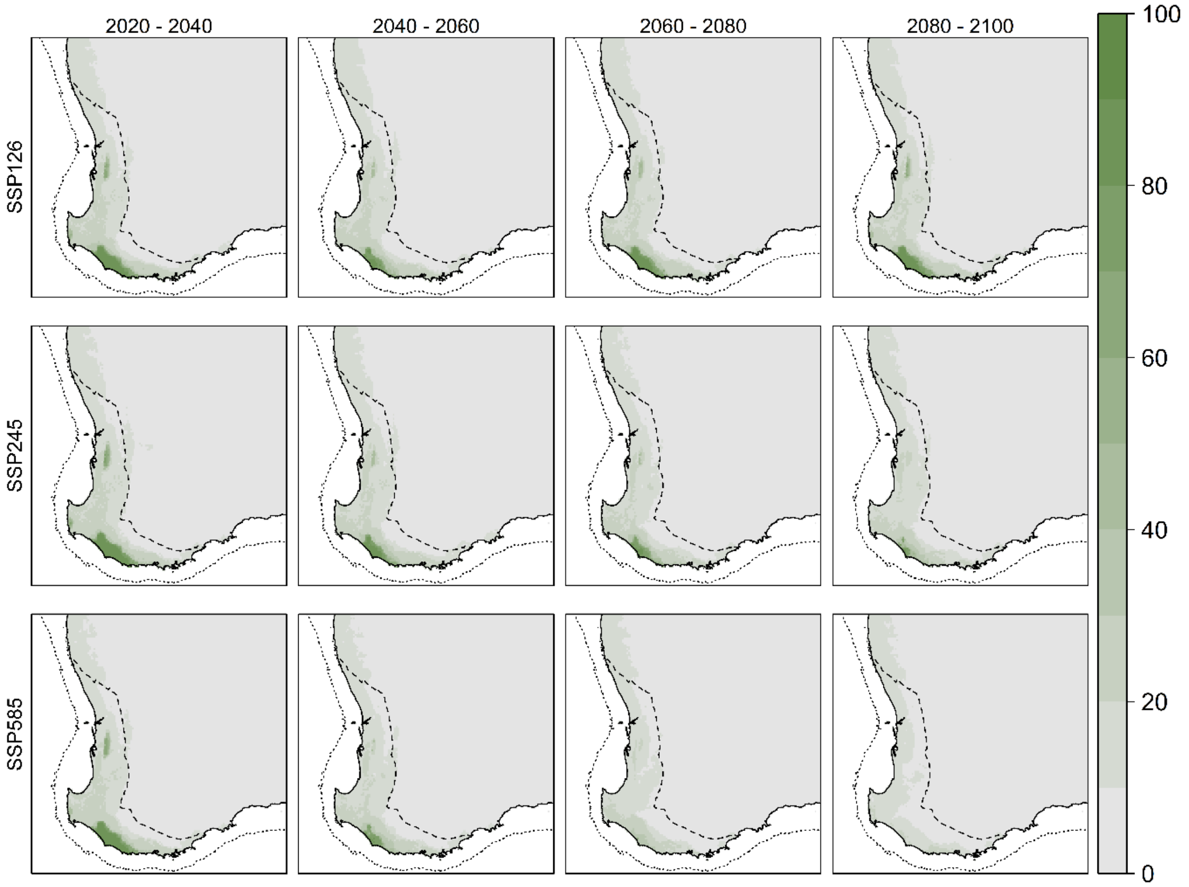


**Figure S11.** Future projections of the LDM for *N. vittata* [B] under climate change. Map grid is separated by climate change scenarios (rows; from least severe at top to most severe at bottom) and time (from left to right).
